# Endothelial cell signature in muscle stem cells validated by VEGFA-FLT1-AKT1 axis promoting survival of muscle stem cell

**DOI:** 10.1101/2021.08.28.458037

**Authors:** Mayank Verma, Yoko Asakura, Xuerui Wang, Kasey Zhou, Mahmut Ünverdi, Allison P. Kann, Robert S. Krauss, Atsushi Asakura

## Abstract

Endothelial and skeletal muscle lineages arise from common embryonic progenitors. Despite their shared developmental origin, adult endothelial cells (ECs) and muscle stem cells (MuSCs) (satellite cells) have been thought to possess distinct gene signatures and signaling pathways. Here we shift this paradigm by uncovering how adult MuSC behavior is affected by the expression of a subset of EC transcripts. We used several computational analyses including single-cell RNAseq to show that MuSCs express low levels of canonical EC markers. We demonstrate that MuSC survival is regulated by one such prototypic endothelial signaling pathway (VEGFA-FLT1). Using pharmacological and genetic gain- and loss-of-function studies, we identify the FLT1-AKT1 axis as the key effector underlying VEGFA-mediated regulation of MuSC survival. All together, our data support that the VEGFA-FLT1-AKT1 pathway promotes MuSC survival during muscle regeneration, and highlights how the minor expression of select transcripts is sufficient for affecting cell behavior.

## Introduction

Skeletal muscle and endothelial cells (ECs) and their progenitors from the trunk and limbs are derived from the somites during early developments. Previous works demonstrated the existence of bipotent progenitors which express both Pax3 and FLK1 (Eichmann et al., 1993; Ema et al., 2006; Esner et al., 2006; Kardon et al., 2002; Tozer et al., 2007). These bipotent progenitors migrate into trunk and limb buds from ventrolateral region of the somites to generate MyoD(+) myogenic cells followed by skeletal muscle and PECAM1(+) ECs followed by vasculatures (Hutcheson and Kardon, 2009; Kardon et al., 2002; Lagha et al., 2009; Mayeuf-Louchart et al., 2014; Mayeuf-Louchart et al., 2016). In addition, FLK1(+) cells give rise to myogenic cells during development and oncologic transformation (Drummond and Hatley, 2018; Mayeuf-Louchart et al., 2014; Motoike et al., 2003). Lastly, multipotent mesoangioblasts, vessel-associated stem cells, have been identified in embryonic dorsal aorta (Minasi et al., 2002). These cells are able to differentiate into several types of mesodermal tissues including skeletal muscle and ECs (Roobrouck et al., 2011). Interestingly, these myogenic cells show the same morphology as muscle satellite cells (MuSCs), stem cell populations for skeletal muscle, and express a number of myogenic and EC markers such as MyoD, M-cadherin, FLK1 and VE-cadherin (De Angelis et al., 1999). However, it is not clear whether adult MuSCs derived from these bipotent progenitors still maintain canonical EC signals. Curiously, blood vessel-associated myoendothelial cell progenitors that express both myogenic and EC markers, and are able to differentiate into myogenic cells following transplantation have been identified in the interstitial spaces of both murine and human adult skeletal muscle (Tamaki et al., 2002; Zheng et al., 2007; Huang et al., 2014). However, the relationship between these myoendothelial cell progenitors and MuSCs remains unclear.

Vascular endothelial growth factor (VEGF), specifically VEGFA modulates many biological aspects including angiogenesis through its two receptors, FLT1 and FLK1. Although FLK1 possesses stronger signaling capability and the major signaling receptor tyrosine kinase (RTK) for VEGFA, FLT1 has considerably higher affinity for VEGF but weaker cytoplasmic signaling capability. In normal tissue, FLT1 acts as a decoy receptor and a sink trap for VEGF thereby preventing excessive normal and pathological angiogenesis. In addition, there are two co-receptors for VEGFA (NRP1 and NRP2) that function with FLK1 to modulate VEGFA signaling. While VEGF signaling has been extensively studied for its role in development, proliferation, and survival of endothelial cells (ECs), its role in non-vascular systems such as neuron and bone has only recently been appreciated (Liu et al., 2012; Okabe et al., 2014; Poesen et al., 2008). Skeletal muscle tissue is the most abundant producer of VEGFA in the body. It has already been extensively studied in the skeletal muscle fibers in models of *Vegfa* knockout mice (Olfert et al., 2009; Tang et al., 2004; Wagner et al., 2006) as well as *Vegfa* overexpression (Arsic et al., 2004; Bouchentouf et al., 2008; Messina et al., 2007; Yan et al., 2005).

Adult skeletal muscle also contains the tissue resident muscle stem cell population, termed MuSCs, which mediate postnatal muscle growth and muscle regeneration (Motohashi and Asakura, 2014). After muscle injury, quiescent MuSCs initiate proliferation to produce myogenic precursor cells, or myoblasts. The myoblasts undergo multiple rounds of cell division before terminal differentiation and formation of multinucleated myotubes by cell fusion. Importantly, the MuSC-derived myoblasts also express VEGFA, which has been shown to increase the proliferation of myoblasts (Christov et al., 2007). Our data obtained from genetical model mice demonstrated that MuSCs express abundant VEGFA, which recruits ECs to establish vascular niche for MuSC self-renewal and maintenance (Verma et al., 2018). In addition, VEGFA and its receptors are expressed in the myoblast cell line, C2C12 cells, and the signaling can induce cell migration and protect apoptotic cell during myogenic differentiation *in vitro* (Bryan et al., 2008; Germani et al., 2003; Mercatelli et al., 2010). However, it is not clear whether MuSCs also express VEGF receptors and if cell-autonomous VEGFA signaling plays an essential roles in MuSC fucction during muscle regeneration *in vivo*.

We have previously shown that *Flt1* heterozygous gene knockout and conditional deletion of *Flt1* in ECs display increased capillary density in skeletal muscle, indicating the essential roles for *Flt1* in adult skeletal muscle. More importantly, when crossed with the Duchenne muscular dystrophy (DMD) model *mdx* mice, these mice show both histological and functional improvements of their dystrophic phenotypes. This was due to the effect of increased ECs leading to an increase in MuSCs (Verma et al., 2010; Verma et al., 2019). However, the effect of VEGFA on MuSC *in vivo* remained unknown. We found that MuSCs express low levels of canonical EC markers including VEGF receptors using single cell transcriptomics. Therefore, we examined the effects of VEGFA on MuSCs and show that it has a drastic effect on cell survival in the via its receptor FLT1 by signaling through AKT1.

## Results

### EC gene signal including *Vegf receptors* in MuSCs

EC signatures in MuSCs has been seen in several gene expression data sets (Figure S1A-D, Table S1) (Fukada et al., 2007; Ryall et al., 2015; van Velthoven et al., 2017; Yue et al., 2020). However, with the lack of EC control, we questioned whether these were true expression or artifact. To isolate EC and MuSC populations, we first crossed the *Flk1^+/GF^*^P^ mice to label the ECs of the vasculature (Ema et al., 2006) and the *Pax7^+/CreERT2^*:*ROSA26^+/Loxp-stop-Loxp-tdTomato^* (*Pax7^+/CreERT2^:R26R^+/tdT^*) mice to mark the MuSC lineage (Murphy et al., 2011; Verma et al., 2018). We performed bulk RNA sequencing (RNAseq) on FACS sorted ECs and MuSCs as well as freshly isolated single muscle fibers (Figure 1A, S1E-G). We found that single muscle fibers routinely have ECs fragments attached to the fiber (Figure S1G) and so we removed such fibers based of Flk1^GFP^ expression from the samples collected for sequencing. We surveyed for canonical genes for each cell type (Figure 1B) and found minimal but reliable expression of canonical ECs genes such as *Pecam1*, *Cdh5*, *Kdr*, and *Flt1* in MuSCs.

**Figure 1.**
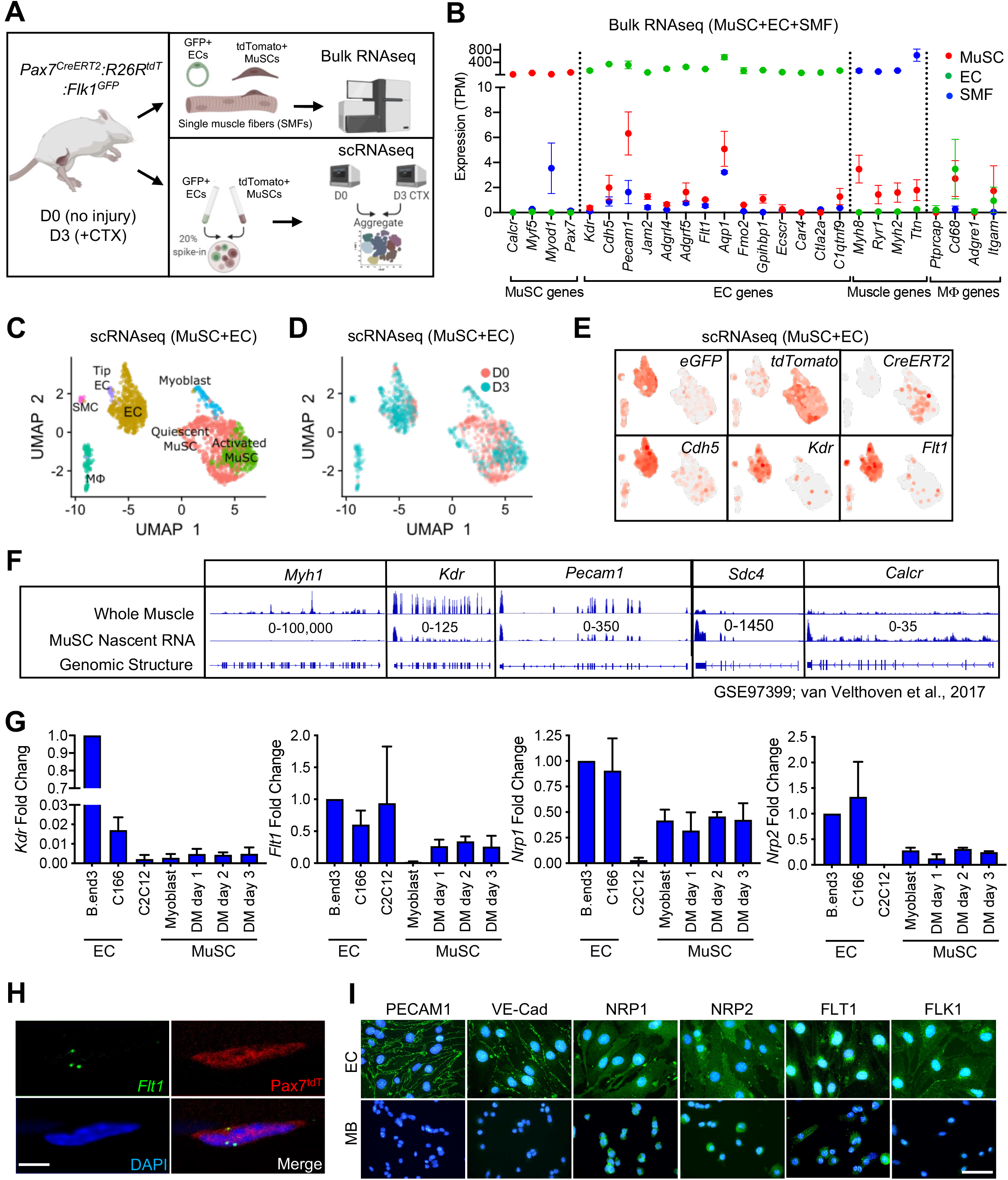
EC gene signal including VEGF receptor genes in MuSCs. A) Experimental schema for bulk and single cell sequencing from the *Pax7^CreERT2^:R26R^tdT^:Flk1^GF^*^P^ mice. Bulk sequencing performed on MuSCs, ECs and single muscle fibers (SMFs) from uninjured muscle. FACS sorted MuSCs and ECs from uninjured and regenerating TA muscle (3 days following CTX) were run separately on the 10X single cell platform and aggregated. B) Bulk RNAseq showing EC signature in MuSCs. Subset dividing genes that are commonly used to delineate cell identity for MuSCs, ECs and SMFs. Last column shows genes that define macrophages (Mφ), which should not be highly expressed in any on our cell types. Red dots indicate MuSCs, green dots indicate ECs and blue dots indicate SMFs. C) UMAP from aggregated single cell RNAseq shows expression of different phases of MuSCs (quiescent MuSCs, activated MuSCs and myoblasts), ECs (tip ECs and ECs) and from likely contaminant cells such as macrophages (Mφ) and smooth muscle cells (SMC). D) UMAP from aggregated data visualized by sample day showing MuSCs segregated by the sample day but overlap in the EC population. Red dots indicate intact (day 0) and blue dots indicate 3 days following CTX. E) Expression of quality control genes such as *eGFP*, *tdTomato*, *CreERT2* and EC genes such as *Cdh5*, *Kdr* and *Flt1*. F) Genome browser tracks of whole muscle and TU-tagged MuSC nascent RNA (GSE97399, van Velthoven et al., 2017). *Kdr* and *Pecam1* expression can be found in the MuSC fraction. As control, *Myh1* is highly expressed in the whole muscle preparation but largely absent in the MuSC fraction. *Sdc4* and *Calcr* are highly expressed in MuSC and less so in the whole muscle fraction. G) qPCR for *Kdr, Flt1, Nrp1* and *Nrp2* in EC lines (B.end3 and C166), muscle cell line (C2C12) and MuSC-derived myoblasts in growth and differentiation medium (DM) shows low level expression of VEGFRs and VEGF co-receptors. H) RNAScope of *Flt1* on freshly isolated single muscle fibers from *Pax7^tdT^* mice shows *Flt1* expression (green) and tdTomato (red) in MuSCs. Nuclei were counterstained with DAPI (blue). Scale bar indicates 5 µm. I) Immunostaining for PECAM1, VE-cadherin (VE-Cad), VEGFA co-receptors (NRP1 and NRP2) and VEGFA receptors (FLT1 and FLK1) in B.End3 EC cell line and MuSC-derived myoblasts (MB). Nuclei were counterstained with DAPI (blue). Scale bar indicates 20 µm.

It is possible that these EC signatures detecting in MuSCs were not due to small amounts of contaminating ECs with very high expression of the canonical EC genes skewing the average expression in MuSC RNA samples. To rule out this possibility, we performed single cell RNAseq (scRNAseq) on MuSCs and ECs isolated from mouse hind limb muscle from both basal condition and 3-days post injury to look at both quiescent and activated MuSCs from the reporter mice specified above (Figure 1A). We could reliably delineate injured and activated MuSCs via side and forward scatter (Figure S1H, S1I). We FACS isolated cells from both days separately and spiked in 20% of the ECs into the MuSCs, and performed scRNAseq for each time point (Figure 1A). We performed sequencing with ∼300K read/cell compared with the commonly used sequencing with 60K reads/cell, in order to maximize the possibility of detecting low abundance transcripts (Zhang et al., 2020). In the aggregated dataset, the MuSCs showed low overlap between D0 and D3 owing to the different stages of the myogenic differentiation cycle, while the ECs clusters showed near perfect overlap (Figure 1C, 1D). While drastic morphological changes in ECs have been shown during muscle regeneration (Hardy et al., 2016), transcriptomic changes are much more tapered, especially compared with MuSCs (Latroche et al., 2017). We were able to deconvolve the quiescent MuSCs from the activated and differentiating MuSCs, ECs, and other cell types from gene signatures. (Figure S1J). We also introduced an artificial chromosome loci in our sequencing reference genome to allow for mapping of custom genes such as *eGFP-SV40* and *tdTomato-WPRE-BGHPolyA* transgenes and were able to confirm high expression of these genes to their respective clusters (Figure 1E). Importantly, data from scRNAseq were able to recapitulate the minimal expression of canonical EC genes in the MuSC clusters such as *Cdh5* (Figure 1E) as seen in our Bulk RNAseq results (Figure 1B). These included the *Vegf receptors Flk1* (*Kdr*) and *Flt1* (Figure 1E).

As a quality control measure, we mapped parts of the three transgene genes, *eGFP* from *Flk1^+/GF^*^P^, and *tdTomato* and *CreERT2* from *Pax7^+/CreERT2^:R26R^+/tdT^* that can be detected in ECs and MuSCs, respectively, as expected (Figure S1L). Surprisingly, we also found *eGFP* in the MuSCs and *tdTomato* in EC fraction, while the *CreERT2* expression remained mainly restricted to the MuSCs (Figure S1K). FACS analysis and FACS-sorted cells confirmed that GFP(+) and tdTomato(+) cells are exclusively restricted as ECs and MuSCs, respectively (Figure S1E, S1F). Therefore, we hypothesized that this was due to the ambient free mRNA from the digested cells that is intrinsic to any droplet based single cell sequencing platform. By using SoupX (Young and Behjati, 2020), we performed careful background subtraction using genes expressed exclusively in myofibers as our negative control and genes validated by *in situ* hybridization as a positive control (Figure S1K) (Kann and Krauss, 2019). We observed decreased but sustained *eGFP* expression in the MuSC fraction and *tdTomato* expression in the EC fraction after SoupX subtraction (Figure S1K). In addition, the EC signatures such as *Cdh5* expression in the MuSC fraction was also sustained. These results conclude that MuSCs contain mRNAs from canonical ECs genes. We showed that the canonical EC genes were broadly expressed in the myogenic cells in our dataset (Figure 1E).

Since detection of rare subpopulation in single cell dataset is a factor of cell numbers, we re-analyzed previously published dataset with 2,232 myogenic cells across different states (Torre et al., 2018; De Micheli et al., 2020). We were able to classify cell as quiescent, proliferative vs. differentiating states based on the expression of *Calcr*, *Cdk1* and *Myog*, respectively (Figure S1L). We noticed that EC prototypic markers such as *Flt1* are broadly expressed with small amounts in MuSCs. Complementary data from different laboratories showed the clear expression of EC prototypic markers such as *Cdh5*, *Flt1* and *Kdr*, using microarrays and Bulk-RNAseq (Figure S1A, S1B; Fukada et al., 2007; Ryall et al., 2015). Recently, RNAseq data from fixed quiescent, early activated and late activated MuSCs show that *Flt1* may be transiently upregulated during the early activation process (Figure S1C) (Yue et al., 2020). To confirm whether the EC gene mRNAs were transcribed from MuSCs, we utilized previously published MuSC nascent RNA transcriptome from TU-tagged samples (Gay et al., 2013; van Velthoven et al., 2017). As expected, *Myh1* was represented in the whole muscle but was absent in the TU-tagged MuSCs (Figure 1F), indicating that the nascent MuSCs were devoid of cellular contamination from other cells in the muscle. Inversely, the nascent MuSC transcript was over-represented for MuSC related genes such as *Calcr* and *Sdc4*. Interestingly, we were able to detect EC genes such as *Kdr* and *Pecam1* in the TU-tagged MuSC samples indicating that they were actively transcribed by MuSCs (Figures 1F, S1D).

We also verified the expression of *Vegfr* genes (*Kdr*, *Flt1*, *Nrp1* and *Nrp2*) in MuSCs using RT-qPCR (Figure 1G). In addition, we verified the expression of *Flt1* by performing *in-situ* hybridization using RNAScope on MuSC on whole muscle fiber, which we currently believe to be the gold standard for expression studies (Figure 1H). Finally, in MuSC-derived myoblasts, NRP1and NRP2 expression was detectable with comparable intensity compared with EC cell line, while FLT1 expression was detectable with lower intensity compared with EC cell line (Figure 1I). By contrast, PECAM1, VE-Cadherin and FLK1 expression, which was clearly detected in EC cell line, was undetectable level in myoblasts. Taken together, these data indicate that there are both transcripts of these EC canonical genes and EC canonical proteins in MuSCs.

### VEGFA induces proliferation and cell survival but not differentiation in myoblasts

Since VEGFRs were expressed in MuSCs in small amounts and their ligand, VEGFA, was highly expressed in MuSCs (Figure 2A; Verma et al., 2018), we wanted to investigate whether there were any biological effects to induction by VEGFA. We found that treatment with VEGFA could increase proliferation of MuSC-derived myoblasts at low dose but inhibit proliferation at high dose of VEGFA, a phenomenon that has been previously described in ECs (Noren et al., 2016) (Figure S2A). We saw no effect on differentiation by VEGFA as evaluated by myosin heavy chain (MyHC) staining, fusion index and RT-qPCR (Figure S2B, S2C, S2D). By contrast, crystal violet staining showed that VEGFA could significantly increase survived cell number of myoblasts following UV-mediated apoptotic cell death induction (Figure S2E, S2F). To investigate apoptosis in detail, we optimized Annexin V assay following thapsigargin-mediated endoplasmic reticulum (ER)-stress (Hirai et al., 2010) so that we could study deviation at ∼ED50 while still performing experiments to remove the confounding variable to proliferation from the experimental setup (Figure S2G and S2H). We had previously shown that MuSCs are the predominant cells that secrete VEGFA in skeletal muscle (Figure 2A; Verma et al., 2018) and while adding exogenous VEGFA did not improve cell survival, blocking VEGFA via a soluble form of FLT1-FC increased the number of apoptotic and necrotic myoblasts *in vitro* (Figure 2C-E).

**Figure 2.**
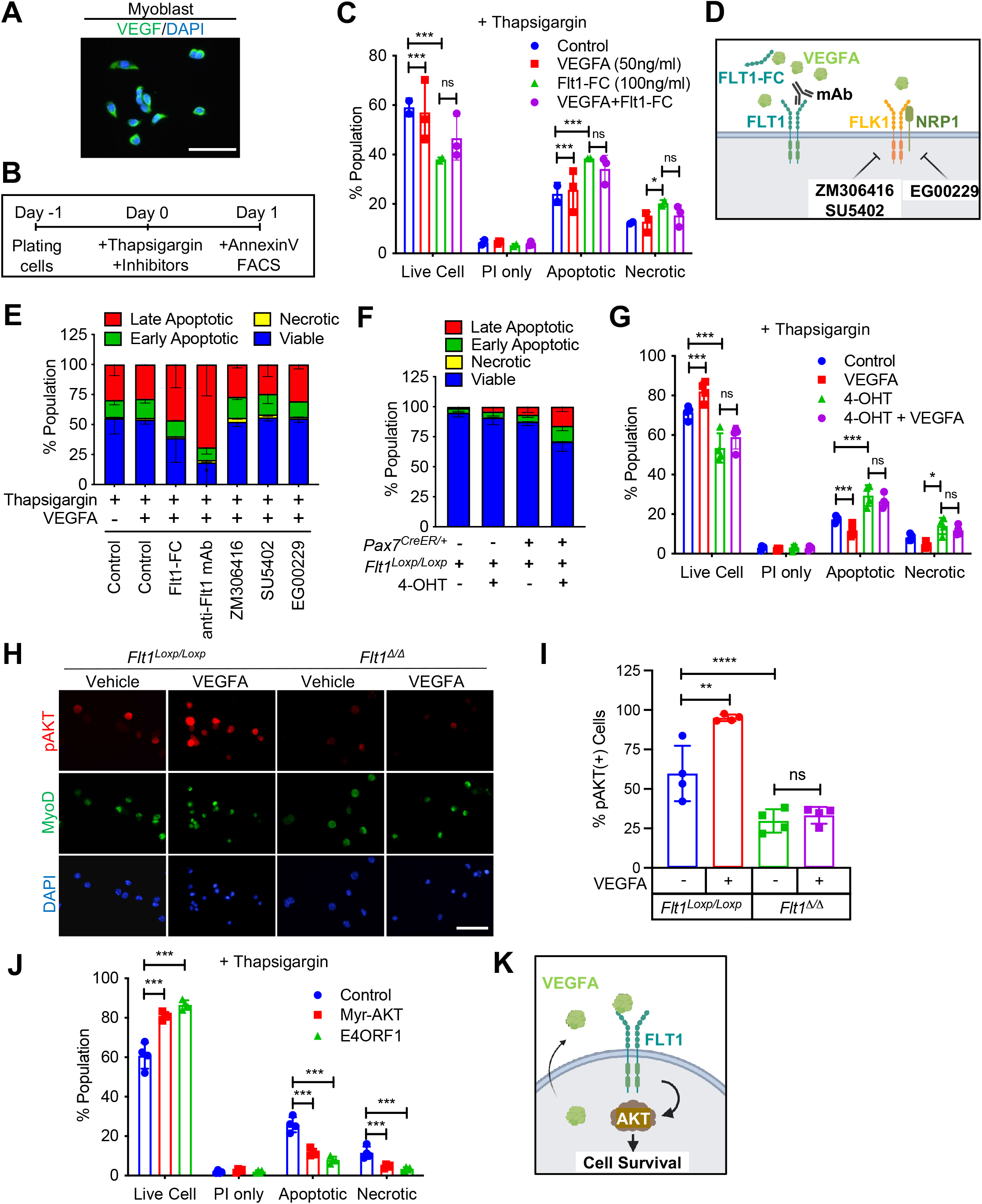
VEGFA-FLT1-AKT1 axis controls apoptosis in MuSC *in vitro*. A) Immunostaining for VEGFA (green) in MuSC-derived myoblasts. Nuclei were counterstained with DAPI (blue). Scale bar indicates 20 µm. B) Experimental scheme for assessing apoptosis following thapsigargin induction in myoblast culture. C) Decreased cell survival in myoblast *in vitro* as VEGFA is blocked using FLT1-FC (a VEFGA trap) following thapsigargin induction. This phenotype is partially rescued with exogenous VEGFA (50 ng/ml). D) Graphical representation of the VEGF pathway inhibitors used in panel E. E) Following thapsigargin induction, apoptotic and necrotic cells are increased with inhibition of FLT1 via FLT1-FC or anti-FLT1 antibody (anti-FLT1 mAb) but not FLK1 (SU5402 and ZM306416) or NRP1-FLK1 inhibition (EG00229) following exogenous VEGFA (50 ng/ml). F) 4-OHT induced deletion of *Flt1* in *Pax7^+/CreER^:Flt1^Loxp/Loxp^* myoblasts is sufficient to reduce cell survival in myoblast without induction of apoptosis. G) Cell survival is decreased *in vitro* in myoblast with thapsigargin induction following 4-OHT mediated deletion of *Flt1* in *Pax7^+/CreER^:Flt1^Loxp/Loxp^* myoblast that is not rescued by exogenous VEGFA. Blue indicates *MuSC-Flt1^+/+^*, red indicates *MuSC-Flt1^+/+^* with 50 ng/ml VEGFA, green indicates *MuSC-Flt1^Δ/Δ^* and purple indicates *MuSC-Flt1^Δ/Δ^* with 50 ng/ml VEGFA. H) Representative images of pAKT (red) in myoblast stained by MyoD (green) in *MuSC-Flt1^+/+^* and *MuSC-Flt1^Δ/Δ^* myoblasts induced with exogenous VEGFA. Nuclei were counterstained with DAPI (blue). Scale bar indicates 50 µm. I) Quantification of pAKT in myoblasts stained by MyoD in *MuSC-Flt1^+/+^* and *MuSC-Flt1^Δ/Δ^* myoblast induced w/wo exogenous VEGFA. VEGFA induction increases pAKT in *MuSC-Flt1^+/+^* myoblasts but this response is lost in *MuSC-Flt1^Δ/Δ^* myoblasts. J) Annexin V quantification of myoblasts transfected with myr-AKT and E4ORF1 to activate AKT1 showed increased cell survival of myoblasts following thapsigargin induction. K) Representative model for VEGFA-FLT1-AKT axis-mediated MuSC survival.

### VEGFA-facilitated cell survival in MuSC-derived myoblasts is mediated through FLT1

To characterize the VEGF receptor responsible for the anti-apoptotic effect of VEGFA on MuSC-derived myoblasts, we used pharmacological inhibitors of the VEGF receptors (Figure 2F). We used blocking antibody for the VEGF receptors FLT1 (anti-FLT1 antibody), small molecule inhibitors for FLK1 (SU4502 and ZM306416) and the FLK1 co-receptor NRP1 (EG00229) following thapsigargin induction (Figure 2D). Surprisingly, inhibiting FLK1, the major signaling RTK for VEGFA, had no effect on myoblasts survival following thapsigargin induction (Figure 3E). By contrast, blocking FLT1 via blocking antibody greatly decreased the survival of myoblasts following thapsigargin induction (Figure 3E). To confirm this interesting result using genetic tools, we obtained myoblasts with *Pax7-CreER*-inducible deletion of *Flt1* mice (*Pax7^+/CreER^:Flt1^Loxp/Loxp^* or *MuSC-Flt1^Δ/Δ^*) and the control mice (*Pax7^+/+^:Flt1^Loxp/Loxp^*). *In vitro* 4-OHT-mediated genetic deletion of *Flt1* (MuSC-*Flt1^Δ/Δ^*) resulted in down-regulation of *Flt1* RNA and FLT1 protein expression (Figure S2I, S2J), and increased spontaneous apoptotic cell death even without induction of apoptosis (Figure 2F). By contrast, *Flt1* deletion did not affect cell proliferation assessed by EdU staining or myogenic differentiation assessed by MyHC staining (Figure S2K-M). When thapsigargin-induced apoptosis was induced, the *MuSC-Flt1^Δ/Δ^* myoblasts had increased apoptosis that was not responsive to exogenous VEGFA (Figure 2G).

**Figure 3.**
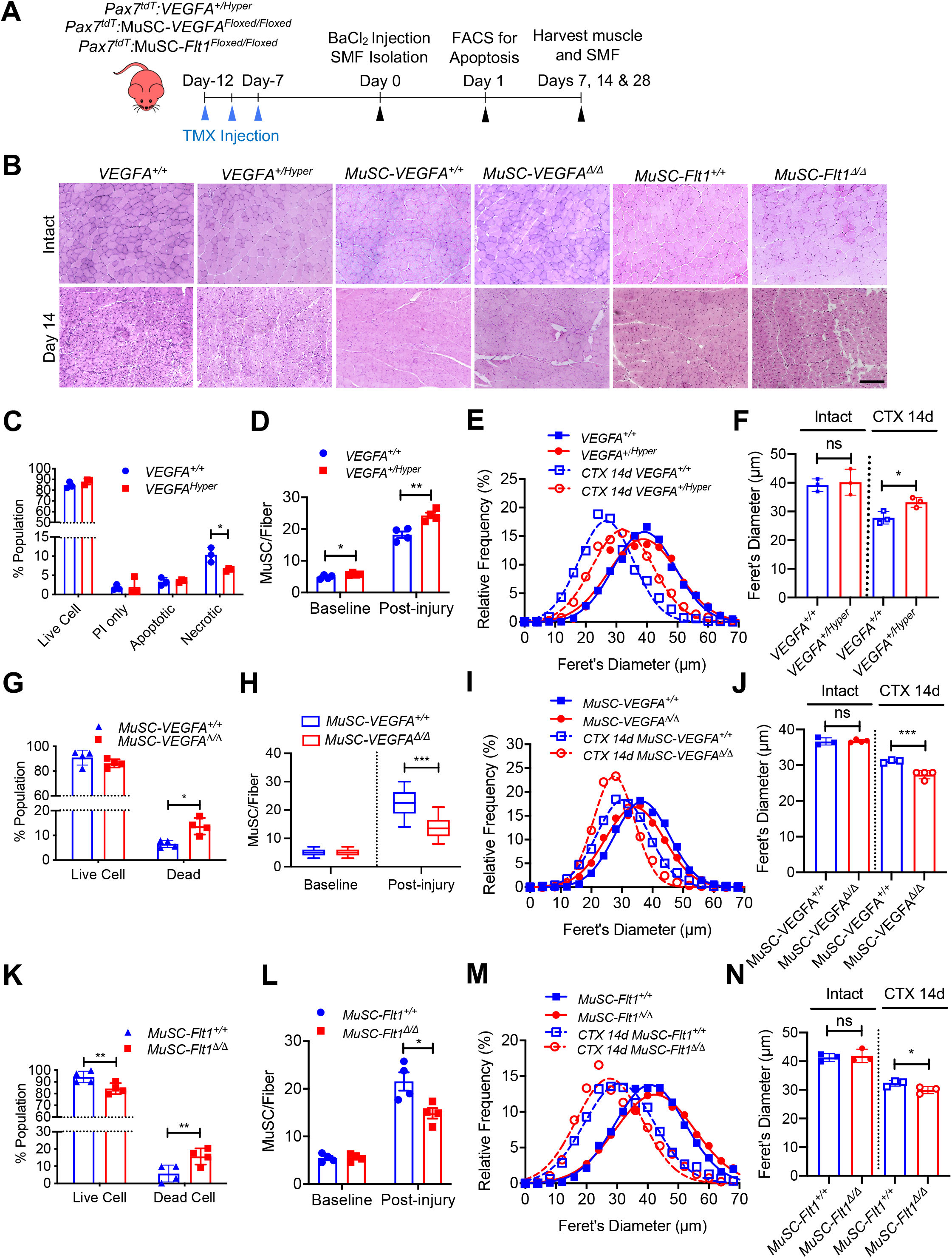
MuSC-derived *VEGFA* and *Flt1* requires proper skeletal muscle. A) Experimental schema detailing the experiments performed in this figure. The *Pax7^+/CreER^:R26R^tdT^:VEGFA^+/Hyper^ (VEGFA^+/Hyper^) Pax7^+/CreER^:R26R^tdT^:VEGFA^Loxp/Loxp^ for MuSC-VEGFA^Δ/Δ^* and *Pax7^tdT^:Flt1^Loxp/Loxp^* for *MuSC-Flt1^Δ/Δ^* lines were pulsed with tamoxifen (TMX) prior to investigation. B) Representative H&E-stained images for intact and on 14-day post injury TA muscle from *MuSC-VEGFA^+/Hyper^, MuSC-Flt1^Δ/Δ^ and MuSC-VEGFA^Δ/Δ^* mice and their representative controls. Scale bar indicates 50 µm. C) Annexin V staining show less necrotic cells in MuSC from *VEGFA^+/Hyper^* mice compared with the control one day following injury. D) Quantification of MuSCs from single muscle fibers show increased MuSCs in *VEGFA^+/Hyper^* EDL muscle compared with the control mice at base line and 14 days post injury. E) Fiber size distribution and F) mean feret’s diameter of uninjured and regenerating muscle 14 days post injury from *VEGFA^+/Hyper^* and control mice show no difference at baseline but an increase in fiber diameter following injury. G) Annexin V staining show increased dead cells in MuSCs from *MuSC-VEGFA^Δ/Δ^* mice one day following CTX compared with the control *MuSC-VEGFA^+/+^* mice. H) Quantification of MuSCs from single muscle fiber at base line and 14 days post injury. I) Fiber size distribution and J) mean feret’s diameter of uninjured and regenerating muscle 14 days post injury from *MuSC-VEGFA^Δ/Δ^* and *MuSC-VEGFA^+/+^* mice show no difference at baseline but a decrease in fiber diameter following injury. K) Annexin V staining show increased apoptosis in MuSCs from *MuSC-Flt1^Δ/Δ^* mice one day following injury compared with the control *MuSC-Flt1^+/+^* mice. L) Quantification of MuSCs from single muscle fiber show decreased MuSCs in *MuSC-Flt1^Δ/Δ^* EDL muscle at base line and 14 days post injury compared with the control *MuSC-Flt1^+/+^* mice. M) Fiber size distribution and N) mean feret’s diameter of uninjured and regenerating muscle from *MuSC-Flt1^Δ/Δ^* and compared with the control *MuSC-Flt1^+/+^* mice show no difference at baseline but a decrease in fiber diameter following injury.

### AKT signaling is involved in apoptosis of muscle stem cells

VEGFA signaling is mediated through Extracellular signal-Regulated Kinase (ERK), p38 Mitogen-Activated Protein Kinase (MAPK), and Protein kinase B (AKT). In ECs, VEGFA is known to protect cells from apoptosis via AKT (Domigan et al., 2015; Lee et al., 2007). However, it is not known whether VEGFA can similarly activate AKT in MuSC-derived myoblasts. While the role of AKT has been explored in proliferation and differentiation in myoblasts, its role in apoptosis has not been well characterized (Loiben et al., 2017). We assessed for AKT activation via phosphorylated AKT (pAKT) in MuSC-derived myoblasts. We found that exogenous VEGFA could induce AKT phosphorylation (pAKT) (Figure 2H, 2I). This response was blunted in *MuSC-Flt1^Δ/Δ^* myoblasts and was no longer responsive to VEGFA (Figure 2H, 2I). Lastly, we wanted to confirm that AKT activation could improve myoblast survival. We infected lentiviral *E4ORF1* or *MyrAKT* vectors in myoblasts (Figure 2J), both of which gene products have been shown to specifically activate AKT without activating ERK or p38 (Kobayashi et al., 2010). We found that overexpression of either of these genes improved cell survival compared with the control *in vitro* following induction of apoptosis via thapsigargin (Figure 2J). These data establish FLT1-AKT as the cascade linking VEGFA to apoptosis in MuSC-derived myoblasts during muscle regeneration (Figure 2K).

### VEGFA-FLT1 pathway protects MuSCs from apoptosis *in vivo*

Endogenous and exogenous VEGFA have been shown to regulate cell survival and protect ECs from apoptosis *in vitro* (Gerber et al., 1998; Lee et al., 2007). To assess whether additional VEGFA had an effect of MuSC behaviors *in vivo*, we used mice carrying the *VEGFA^+/Hyper^* allele for injury-mediated tibialis anterior (TA) muscle regeneration following BaCl_2_ injection (Figure 3A, 3B) (Miquerol et al., 1999). MuSC-derived myoblasts from *Pax7^+/tdT^:VEGFA^+/Hyper^* mice showed around 2.8 fold increased expression of *Vegfa* but not the *Vegfrs* compared with myoblasts from wild-type mice (Figure S3A). Interestingly, while treatment with VEGFA alone had no effect on apoptosis *in vitro*, the MuSCs from *Pax7^+/tdT^:VEGFA^+/Hyper^* mice showed decreased cell death in regenerating muscle by 1 day following BaCl_2_ injection (Figure 3C). Consequently, single muscle fibers from *Pax7^+/tdT^:VEGFA^+/Hyper^* mice showed increased number of MuSCs, compared with those from *Pax7^+/tdT^:VEGFA^+/+^* mice by 28 days following BaCl_2_ injection (Figure 3D). In addition, muscle regeneration was promoted in *VEGFA^+/Hyper^* mice in the early and late muscle repair processes as judged by fiber diameter (Figures 3B, 3E, 3F, S3B-D).

We then performed the reciprocal experiment to investigate the consequence of *Vegfa* loss in MuSCs *in vivo*, and utilized MuSC-specific *Vegfa* knockout mice (*Pax7^+/CreER^:VEGFA^Loxp/Loxp^*). We have previously shown that vasculature in the MuSC-*VEGFA^Δ/Δ^* mouse muscle is perturbed and the proximity between the MuSC and EC is increased (Verma et al., 2018). However, the functional consequences of this remained unknown. We confirmed that clear downregulatin of VEGFA protein in MuSC-derived myoblasts isolated from MuSC-*VEGFA^Δ/Δ^* mice (Figure S3E). We noticed that deletion of *Vegfa* in MuSCs in the *MuSC-VEGFA^Δ/Δ^* mouse muscle led to an increase in the proportion of dead MuSCs following BaCl_2_ injection (Figure 3G). Consequently, the number of MuSCs in the MuSC-*VEGFA^Δ/Δ^* muscle were significantly reduced following recovery after injury (Figure 3H). There was no difference in the MuSC numbers in MuSC-*VEGFA^Δ/Δ^* muscle at homeostasis. In addition, the muscle had a regenerative defect as indicated by the shift in fiber size distribution and increased adipose following muscle injury (Figure 3B, 3I, 3J, S3F-H). While a limitation of this experiment is that the MuSC fusion into the fiber also deletes *Vegfa* from the fiber themselves, muscle fiber specific deletion of *Vegfa* has not shown to have an effect on fiber size (Delavar et al., 2014). These data indicate that cell intrinsic VEGFA improves cell survival of MuSCs and that loss of MuSC-derived VEGFA results in reduced muscle regeneration.

Since FLT1 but not FLK1 was detected in MuSCs and MuSC-derived myoblasts, we asked whether the *Flt1* had an effect on MuSC survival *in vivo,* we evaluated cell death in MuSCs from *MuSC-Flt1^Δ/Δ^* mouse muscle. We induced muscle regeneration using BaCl_2_ for 1 day and assessed for cell death in MuSCs. As seen *in vitro*, we found that loss of *Flt1* in MuSCs (Figure S2I, S2J) resulted in increased cell death during early regeneration (Figure 3K). Consequently, single muscle fibers from *MuSC-Flt1^Δ/Δ^* mice showed decreased number of MuSCs, compared with those from *MuSC-Flt1^+/+^* mice by 28 days following BaCl_2_ injection (Figure 3L). We also examined the long-term *in vivo* consequence of deleting *Flt1* from MuSC. There was no significant muscle phenotype in MuSC-*Flt1^Δ/Δ^* muscle at homeostasis (Figures 3B, 3M, 3N). However, following injury, the MuSC-*Flt1^Δ/Δ^* muscle had a modest regenerative defect as indicated by the shift in fiber size distribution following muscle injury (Figures 3B, 3M, 3N, S3J-N).

### VEGFA-FLT1 pathway regulates muscle pathology in DMD model mice

While angiogenic defects have been reported in the *mdx* mice as well as in golden retrieval muscular dystrophy (GRMD; canine model of DMD) (Latroche et al., 2015; Verma et al., 2019, 2010; Kodippili et al., 2021; Podkalicka et al., 2021), it is not clear whether VEGF family and its receptors are implicated in human dystrophinopathies. We probed the VEGF ligands and receptors in microarrays (Table S1) from skeletal muscles and MuSCs from *mdx* mice (Pallafacchina et al., 2010; Tseng et al., 2002) and skeletal muscles from the *GRMD* (Vieira et al., 2015). *Vegfa* was downregulated in both models (Figure S4A). *Flt1* was also downregulated in *GRMD* but not *mdx* muscles. To examine whether VEGF signaling is altered in DMD patients, we performed gene expression analysis on previously available data from microarrays from patients with DMD (Chen et al., 2000). We also aggregated and probed microarray data from muscle biopsies of patients with various neuromuscular diseases or of healthy individuals after exercise (Bakay et al., 2006). In the microarray data, *Vegfa* expression was increased after an acute bout of exercise, and *Vegfa* expression was reduced in ALS muscle, BMD muscle, as well as both early and late phases of DMD muscle (Figure S4A). These data indicate that *Vegfa* expression is decreased in dystrophinopathy, and thus may benefit people with DMD by increasing VEGFA as a therapeutic target.

Therefore, we crossed the *MuSC-Flt1^Δ/Δ^* mice with the chronically regenerating DMD model mice (*mdx*) to generate *mdx:MuSC-Flt1^Δ/Δ^* mice, and analyzed long term effects of *Flt1* deletion (Figure 4A, S4B). Importantly, we found a significant decrease in fiber diameter and increased fibrosis (Figures 4B-D, S4C) in TA muscle accompanied by a physiological decease in muscle perfusion as shown by laser Doppler flow at 12 months age (Figure 4E) as well as a functional decline in muscle strength as judged by grip strength both acutely and chronically (Figure 4F).

**Figure 4.**
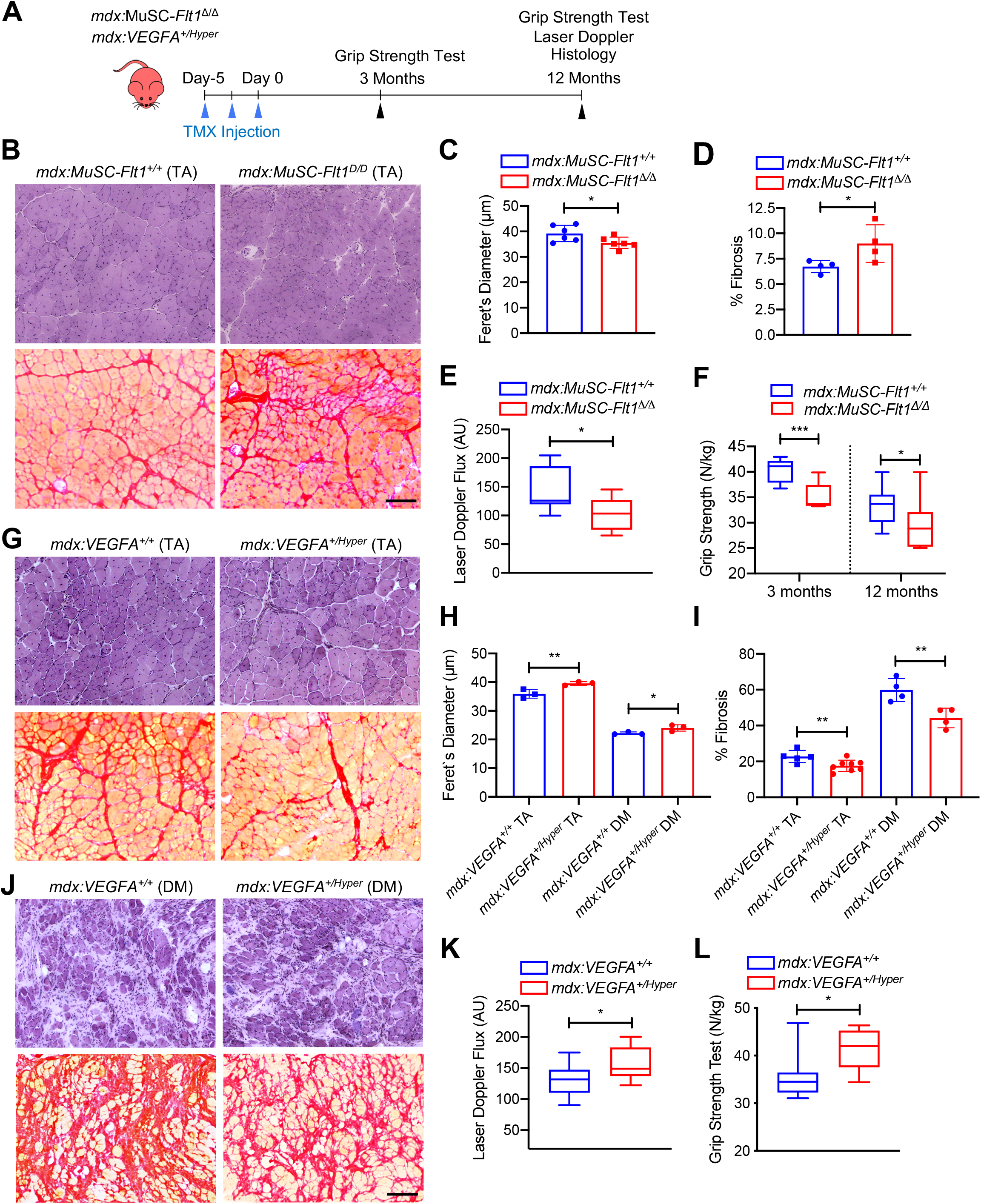
VEGFA-FLT1 pathway in MuSCs regulates muscle pathology in DMD model mice. A) Experimental schema detailing the experiments performed in this figure. The *mdx:Pax7^tdT^:Flt1^Loxp/Loxp^* was pulsed with tamoxifen (TMX) to generate *mdx:MuSC-Flt1^Δ/Δ^* mice prior to investigation. *mdx*:*VEGFA^+/Hyper^* mouse line was used without any induction. B) Representative H&E and Sirus red stain from *mdx:MuSC-Flt1^+/+^* and *mdx:MuSC-Flt1^Δ/Δ^* mouse TA muscle at 3 months of age. Scale bar indicates 50 µm. C) Smaller average fiber size in *mdx:MuSC-Flt1^Δ/Δ^* compared with the control *mdx:MuSC-Flt1^+/+^* mouse TA muscle. D) Increased fibrotic area in *mdx:MuSC-Flt1^Δ/Δ^* compared with the control *mdx:MuSC-Flt1^+/+^* mouse TA muscle. E) Decreased muscle perfusion in *mdx:MuSC-Flt1^Δ/Δ^* compared with the control *mdx:MuSC-Flt1^+/+^* mouse TA muscle. F) Decreased grip strength normalized to body weight in *mdx:MuSC-Flt1^Δ/Δ^* compared with the control *mdx:MuSC-Flt1^+/+^* mouse TA muscle at both 3 and 12 months of age. G) Representative H&E and Sirus red stain from *mdx*:*VEGFA^+/Hyper^* and *mdx*:*VEGFA^+/+^* mouse TA muscle at 3 months from TA muscle (top four panels) and diaphragm muscle (bottom four panels). H) Increased average fiber size in *mdx*:*VEGFA^+/Hyper^* compared with the control *mdx*:*VEGFA^+/+^* mouse TA and diaphragm (DM) muscle. Scale bar indicates 50 µm. I) Decreased fibrosis in *mdx*:*VEGFA^+/Hyper^* compared with the control *mdx*:*VEGFA^+/+^* mouse TA muscle. J) Decreased fibrosis in *mdx*:*VEGFA^+/Hyper^* compared with the control *mdx*:*VEGFA^+/+^* mouse diaphragm (DM) muscle. K) Muscle perfusion is increased in *mdx*:*VEGFA^+/Hyper^* compared with the control *mdx*:*VEGFA^+/+^* mouse TA muscle. L) Grip strength normalized to body weight is increased in *mdx*:*VEGFA^+/Hyper^* compared with the control *mdx*:*VEGFA^+/+^* mice.

By contrast, when we crossed the *VEGFA^+/Hyper^* mice with *mdx* mice (Figure 4A, S4D), we noticed a significant increase in fiber diameter and decreased fibrosis (Figures 4G-J, S4E) in both TA and diaphragm muscle of *mdx:VEGFA^+/Hyper^* mice accompanied by a physiological increase in muscle perfusion as shown by laser Doppler flow at 12 months age (Figure 4K) as well as a functional increase in muscle strength as judged by grip strength (Figure 4L). These data indicate that VEGFA-FLT1 axis is a therapeutic target for the pathology seen in the *mdx* mice

## Discussion

In this report, we performed bulk and single cell RNA sequencing on MuSCs and ECs. Since deep reads can significantly reduce the effect of the technical noise in scRNAseq, it can improve estimation of minor transcriptional state of a given cell (Zhang, 2020). Unexpectedly, we found that MuSCs broadly express EC prototypic markers in small amounts and used multiple different bioinformatics techniques to validate the results. While similar phenomenon in myogenic cells during development and existence of blood vessel-associated myoendothelial cells in the adult skeletal muscle have been previously described, no functional follow up as been performed leading to the questions whether these minor expression profiles were artifacts or functional (De Angelis et al., 1999; Minasi, 2002; Tamaki et al., 2002; Zheng et al., 2007; Roobrouck, 2011; Huang et al., 2014; Charville et al., 2015; Goel et al., 2017; Giordani et al., 2018). Our goal was to see whether this small expression pattern had biological consequences. We ultimately decided to use *Flt1* for further investigations and used RNAscope and immunostaining to validate its expression in MuSCs. We found that *Flt1* indeed exerts a biological function even at a low expression. Signaling through VEGFA-FLT1-AKT can improve cell survival in MuSCs both *in vivo* and *in vitro*.

On a grander scale, our finding of EC prototype markers expressed in MuSC calls into two questions 1) the genes that we used to specify cellular identities and 2) the cellular identity of MuSCs and ECs. The former is important as when we experimentally label, induce or perform Cre-mediated gene knockout experiments based on our assumptions of different gene expression results which may be confounded for these low expressing genes. For example, we have previously investigated both *Flt1* and *Kdr* in mouse muscle using three different reporters and found them to be negative in MuSCs, thereby disregarding their cell-autonomous effect when evaluating global knockouts (Verma et al., 2018, 2010). It is also possible that EC mRNAs are results of transcription from the cell or a result of mRNA transfer from neighboring cells (Desrochers et al., 2016). Of note, the transmission of *tdTomato* mRNA and protein from *Pax7^+/CreERT2^:R26R^+/tdT^* mice used in this study has been recently shown via exosome, opening up the possibility of transmission of other mRNA from MuSC to ECs (Murach et al., 2020). The later is an interesting phenomenon form a developmental point of view. MuSCs and ECs arise from a bipotent progenitor originated from somites during early development (Kardon et al., 2002; Hutcheson, 2009; Lagha, 2009; Mayeuf-Louchart, 2014; Mayeuf-Louchart, 2016). Therefore, it is possible that there is a permissive chromatin state that allows for expression of reciprocal genes in the two populations. Along the lines of these observations, FLK1(+) or VE-cadherin(+) cells can contribute to myogenic cells *in vitro* and after cell transplantation (Tamaki et al., 2002; Le Grand et al., 2004; Zheng et al., 2007; Huang et al., 2014;), and during development (Drummond and Hatley, 2018; Mayeuf-Louchart et al., 2014; Motoike et al., 2003). Important notion is that the PDGFRα(-)FLK1(+) population exhibited restricted potential to differentiate into the MuSCs in injured muscle (Sakurai et al., 2008). Interestingly, in the zebrafish, exogenous expression of *Etv2* in the fast muscle can lead to transdifferentiation of muscle fibers into functional vessels so there is evidence of cell fate flexibility (Veldman et al., 2013). The potential of EC transdifferentiation was also examined by ETV2 overexpression in five human cell types, skeletal muscle cells, adipose-derived mesenchymal stem cells, umbilical cord-derived mesenchymal stem cells, embryonic lung fibroblast cells and skin fibroblast cells. Among them, human skeletal muscle cells showed the highest amenability for this EC induction following infection with ETV2 lentivirus vector (Yan et al., 2019). Conversely, *Etv2*-deficient vascular progenitors can differentiate into skeletal muscle cells (Chestnut et al., 2020). It would be interesting to see whether other EC gene signatures also have functional consequences in the MuSC or muscle at large.

We decided to focus on function of *Flt1* among several EC genes expressed in MuSCs for further investigations on MuSC biology. Our pharmacological and genetic analyses demonstrate that MuSC-derived VEGFA has a drastic effect on cell survival in the via its receptor FLT1 by signaling through AKT1. While VEGFA binds to both FLT1 and FLK1, VEGFB and PGF only bind to FLT1. This creates a scenario where PGF and VEGFB binding can sequester FLT1, increasing free VEGFA availability for VEGFA-FLK1 binding which is the major VEGF signaling pathway for many cell types (Vempati et al., 2014). While PGF is not normally expressed in adult tissues, VEGFB is expressed in the MuSCs and muscle fiber (data not shown). Importantly, the VEGFB-FLT1 axis has also been shown to inhibit apoptosis in retina and brain cells in mouse models of ocular neurodegeneration and stroke (Li et al., 2008). While our results cannot rule out the involvement of VEGFB in protection of MuSC apoptosis, we provide evidence from both pharmacological and genetic data to indicate that VEGFA is involved.

Despite drastic effect of VEGFA-FLT1 on apoptosis *in vitro*, the long-term consequences of *in vivo* deletion of *Flt1* in the MuSC compartment were modest compared with deletion of *Vegfa* in the MuSCs unless crossing with *mdx* mice. Although *Vegfa* is required for both MuSC survival and recruitment of vascular niche (Verma et al., 2018), in the steady state, the MuSC turnover may be low enough that the apoptotic stress burden is low. VEGFA improves cell survival during the proliferative stage following injury, however this transient improvement in survival has only a modest impact on the final regenerative process as shown in *mdx:VEGFA^+/Hyper^* mice.

VEGFA and FLT1 targeted therapies are being explored as both pro- and anti-angiogenic therapies for several indications including retinal degeneration, cancer, pre-eclampsia and neuromuscular diseases (Bae et al., 2005; Keifer et al., 2014; Mac Gabhann et al., 2011; Verma et al., 2010; Verma et al., 2019; Bosco et al., 2021; Xin et al., 2021). As these therapies mature, it will be important to ascertain the MuSC-specific effects of VEGFA and FLT1.

## Materials and Methods

### Mice

*Flt1^LoxP/LoxP^* were obtained from Gua-Hua Fong (Ho et al., 2012). *B6.Cg-Pax7^tm1(cre/ERT2)Gaka/J^* (*Pax7^+/CreERT2^*; JAX stock# 017763; Murphy et al., 2011), *B6.Cg-Gt(ROSA)^26Sortm9(CAG-tdTomato)Hze/J^* (*Ai9*; JAX stock # 007909; Madisen et al., 2010), *VEGFA^+/Hyper^* (*Vegfatm1.1Nagy/J*; JAX stock# 027314; Miquerol et al., 1999) and *B6Ros.Cg-Dmd^mdx-5cv^/J* (*mdx^5cv^*; JAX stock #002379; Chapman et al., 1989) were obtained from Jackson Laboratory. *Kdr^tm2.1Jrt/J^* (*Flk1^+/GFP^*) were obtained from Masatsugu Ema (Ema et al., 2006). *B6.Cg-Pax7^tm1(cre/ERT2)Gaka/J^* (*Pax7^+/CreERT2^*) mice were crossed with the *B6.Cg-Gt(ROSA)^26Sortm9(CAG-tdTomato)Hze/J^* (*Ai9*) to yield the *Pax7^+/CreERT2^:R26R^tdT^*(*Pax7^tdT^*) mice. *Pax7^tdT^* mice were bred with the *VEGFA^+/Hyper^* and *Flk1^+/GFP^* to obtain *Pax7^+/tdT^*:*VEGFA^+/Hyper^* and *Pax7^+/tdT^:Flk1^+/GFP^* mice. *VEGFA^LoxP/LoxP^* mice obtained from Napoleone Ferrara (Gerber et al., 1999) were crossed with *Pax7^+/CreERT2^* to yield the *Pax7^+/CreERT^: VEGFA^LoxP/LoxP^* mice. *Flt1^LoxP/LoxP^* mice obtained from Guo-Hua Fong (Ho et al., 2012) were crossed with *Pax7^+/CreERT2^* to yield the *Pax7^+/CreERT^:Flt1^LoxP/LoxP^* mice. Colonies for all the mice were established in the laboratory. Cre recombination was induced using tamoxifen (Sigma-Aldrich, T5648) dosed as 75 mg/kg body weight x 3 times over one week at 3-6 weeks of age. Mice carrying the wild-type *CreERT2* allele were used for control experiments. TA muscle regeneration was induced by intramuscular injection of 20 µl of 1% BaCl_2_ (Sigma-Aldrich, 342920) or 20 µl of 10 µM Cardiotoxin (CTX) (Sigma-Aldrich, V9125). Mice used for this study is summeried in Table S2.

Genotyping to detect the transgenic and mutant alleles was performed by PCR using the primers described on the web site of Jackson Laboratory shown in Table S3. All primers were synthesized as custom DNA oligos from Integrated DNA technologies (IDT). Genotyping to detect the mutated allele of *mdx^5cv^* was performed by PCR using the primers (0981 and 0982) shown in Table S1. The PCR product DNA was digested with *Dra*III restriction enzyme (New England Biolabs, R3510S). Wild-type allele generated 180 bp and mutant allele generated 50 and 130 bp bands.

The animals were housed in an SPF environment and were monitored by the Research Animal Resources (RAR) of the University of Minnesota. All protocols were approved by the Institutional Animal Care and Usage Committee (IACUC) of the University of Minnesota and complied with the NIH guidelines for the use of animals in research.

### Cell isolation by FACS

*Pax7^tdT^:Flk1^GFP^* mice were utilized for FACS-mediated MuSC and EC isolation as previously described (Asakura et al, 2002; Verma, 2018). We performed extensive validation of the fluorescent reporter mice as previously described (Figure S1A-C; Verma, 2018). Briefly, quiescnet MuSCs and ECs were isolated from the hind limb skeletal muscle of 1–2-mo-old *Pax7^tdT^:Flk1^GFP^* mice after digestion with collagenase type II. FACS was performed on an FACS sorter (BD FACSAria) and data were analyzed using FlowJo (BD Biosciences). Sorting gates, tdTomato(+) for MuSCs and GFP(+) for ECs, were strictly defined based on control cells isolated from wild-type mice and the forward scatter and side scatter gating. Sorted cells were immediately characterized by immunostaining on slide glasses, utilzed for RNA preparation or cultured on collagen-coated plates in the myoblast growth medium as below to obtain MuSC-derived myoblasts and ECs. FACS analysis was performed as previously described (Turaç et al., 2013). Cells were either trypsinized (cultured cells) or a single cell suspension was obtained following enzymatic digestion as whole muscle-derived cells (Asakura, 2002). Cells were then washed with FACS buffer (2% BSA and 1 mM EDTA in PBS) followed by live/dead staining using ZombieNIR (Biolegends, 423105). Cells were washed, then immunostained for cell surface markers. Blocking cells was performed with 1% BSA/PBS, and cells were incubated in fluorescently-conjugated antibody. FACS was performed on a Fortessa X-20 (BD Biosciences) with a 355 nm, 405 nm, 488 nm, 561 nm, and 640 nm lasers.

### Cell culture

MuSC-derived myoblast isolation from adult mice was performed as previously described (Asakura et al, 2001; Motohashi et al., 2014). Briefly, after collagenase type II (Worthington, CLS-2) treatment, dissociated cells from mouse hindlimb muscle were incubated with anti-CD31-PE (eBiosciences, 12-0311), anti-CD45-PE (eBiosciences, 30-F11), anti-Sca1-PE (eBiosciences, Dec-81) and anti-Integrin α7 (MBL International, ABIN487462), followed by anti-PE microbeads (Miltenyi Biotec, 130-048-801), and then performed LD column (Miltenyi Biotec, 130-042-901) separation. Negative cell populations will be incubated with anti-Mouse IgG beads (Miltenyi Biotec, 130-048-402), and then MS coulumn (Miltenyi Biotec, 130-042-201) separation was performed to isolate Integrin α7(+) MuSCs. MuSC-derived myoblasts were maintained in culture on collagen coated plates in myoblast medium containing 20% FBS and 20 ng/ml bFGF (Invitrogen, PHG0263) in HAM’s-F10 medium. Cell cultures were maintained in a humidified incubator at 37°C with 5% CO_2_ and 5% O_2_. 4-hydroxy tamoxifen (4-OHT, Sigma-Aldrich, H6278) treatment (1 µM in EtOH) was used to induce *Flt1* deletion in myoblasts isolated from *Flt1^LoxP/LoxP^:Pax7^CreERT2^* mice. For cell survival assay, 1 x 10^5^ cells were allowed to adhere for one day and starved overnight in 0.1% FBS in HAM’s F10 medium. Then, cells were exposed to 1 µM EdU along with or withpout 2-100 ng/ml recombinant VEGFA (R&D Systems, 493-MV) for 8 hours before being fixed and stained by the Click-iT EdU Alexa Fluor 488 Imaging Kit (Thermo Scientific, C10337). For induction of apoptosis in myoblasts, (1-2 x 10^5^) cells were allowed to adhere to the plates for 16 hours. Thapsigargin-mediated apoptosis was induced by 1 µM of thapsigargin (Sigma-Aldrich, T9033) dissolved in EtOH with or without VEGFA, 100 ng/ml recombinant FLT1-FC (R&D Systems, 7756-FL), 1 µg/ml anti-FLT1 monoclonal antibody (Angio-Proteomie, MAB7072), inhibitors of FLK1, 3 µM ZM306416 (R&D Systems, 2499/1) and 10 µM of SU5402 (R&D Systems, 3300/1) and an inhibitor of NRP1, 30 µM of EG00229 (R&D Systems, 6986/10) for 24 hours. UV light-mediated apoptosis was induced by exposing the cells to UV light in cell culture hood for 45 seconds without medium. After UV exposure, cell survival was assessed 24 hours following culture in 0.1% FBS in HAM’s F10 medium with or without VEGFA using the Crystal violet Assay Kit (Abcam, ab232855) and quantated the Crystal violet dye after solubilization by absorbance at 570 nm. To induce differentiation of myoblasts, the myoblast medium was replaced with differentiation medium that contained DMEM supplemented with 5% horse serum with or withour VEGFA or bFGF for 1 or 3 days followed by anti-sarcomeric myosin heavy chain antibody (Developmental Study Hybridoma Bank, MF-20).

### AKT induction

The lentiviral pCCL-E4ORF1 and pCCL-myrAkt1 constructs were a kind gift from Dr. Jason Butler (Kobayashi et al., 2010). 293FT cells (Thermo Fisher Scientific, R70007) were seeded in DMEM with 10% FBS and transfected with the lentivirus vectors along with pCMV-VSV-G (Addgene, 8454), pRSV-Rev (Addgene, 12253), and pMDLg/pRRE (Addgene, 12251) using PolyJet transfection reagent (Signagen Laboratories, SL100688). The culture supernatant of the transfected 293FT cells was added to MuSC-derived myoblast culture with 0.8 μg/ml polybrene (MilliporeSigma, H9268). pAKT1(+) cells were stainined with anti-pAKT antibody (Cell Signaling, 4060).

### Apoptosis assay

Apoptosis was measured using measured using Annexin V-Biotin Apoptosis Detection Kit (eBioscience™, BMS500BT-100) as per the manufacture’s instruction. Streptavidin-conjugated AlexaFluro-488 was used for detection. Propidium Iodide (PI) was used in all assays except when Pax7tdT(+) cells were utilized or when ZombieNIR (Biolegends, 423105) was used. FACS was performed on a Fortessa X-20 (BD Biosciences) equipped with a 355 nm, 405 nm, 488 nm, 561 nm, and 640 nm lasers.

### Immunostaining of cells

Immunostaining for PECAM1, VE-Cadherin, VEGFA, VEGFRs was performed on collagen coated coverslips. Other immunostaining was performed on 30 mm tissue culture plates. Cells were fixed with 2% PFA for 5 minutes and immunostained as previously described (Verma et al., 2010). For membrane receptor staining, cells were permeabilized with 0.01% saponin (ThermoFisher Scientific, ICN10285525) which was kept in the staining solution until the primary antibodies were washed off. At which time, 0.01% Triton-X was added to all the buffers. The antibodies used for this study are listed in Table S4.

### Single muscle fiber isolation and staining

Extensor digitorum longus (EDL) muscle was dissected and digested with 0.2% collagenase type I (Sigma-Aldrich, C0130) for single muscle fiber isolation as previously described (Verma et al., 2010). Single muscle fibers were fixed with 2% PFA/PBS, permeabilized with 0.2% Triton-X100 and counterstained with DAPI. Anti-Pax7 antibody(+) or tdTomato(+) MuSCs per single muscle fiber were counted manually.

### RNAscope

RNAscope for *Flt1* transcripts was performed as previously described (Kann and Krauss, 2019) on single muscle fibers from *Pax7^tdT^* mice using the RNAscope Probe - Mm-Flt1 (C1) (ACDBio, 415541). Briefly, isolated EDL fibers are fixed in 4% PFA, washed with PBS, and dehydrated in 100% methanol. Subsequently, fibers are rehydrated in a stepwise gradient of decreasing methanol concentrations in PBS/0.1% Tween-20. Fibers are treated with a proteinase for 10 minutes, followed by hybridization, amplification, and fluorophore conjugation steps.

### Histology

The mouse tibialis anterior (TA) muscle was used for all histological analysis. Tissues were frozen fresh using LiN_2_ chilled isopentane and stored at −80°C. Eight μm thick transverse cryosections were used for all histological analysis. Hematoxylin & Eosin (HE) staining were performed as previously described (Verma et al., 2010). Sirius red (Direct Red 80, Sigma-Aldrich, 365548) staining was performed for muscle sections for fibrosis as previously described (Shimizu-Motohashi et al., 2015). Muscle sections were stained in Oil Red O solution (Sigma-Aldrich, O1391-250ML) as previously described (Wang et al., 2017). Microscopic images were captured by a DP-1 digital camera attached to BX51 fluorescence microscope with 10x or 40×UPlanFLN objectives with cellSens Entry 1.11 (all from Olympus). Photoshop (Adobe) and Fiji (NIH) were used for image processing and manually enumerating the fiber diameter (Schindelin et al., 2012).

### Grip strength test

Forelimb grip strength test was performed following a previously published procedure (Aartsma-Rus and van Putten, 2014). Briefly, mice were gently pulled by the tail after fore limb-grasping a metal bar attached to a force transducer (Grip Strength Meter, Columbus Instruments, 1027CSM-D52). Grip strength tests were performed by the same blinded examiner. Five consecutive grip strength tests were recorded, and then mice were returned to the cage for a resting period of 20 minutes. Then, three series of pulls were performed each followed by 20 min resting period. The average of the three highest values out of the 15 values collected was normalized to the body weight for comparison.

### Muscle perfusion

RBC flux was evaluated using the moorLabTM laser Doppler flow meter as previously described (Verma, 2010) with the MP7a probe that allows for collecting light from a deeper tissue level than standard probes according to the manufacturer’s instructions (Moor Instruments). The fur from the right hind leg was removed using a chemical depilatory. Readings were taken using the probe from at least 10 different spots on the TA muscle. The AU was determined as the average AU value during a plateau phase of each measurement.

### RNA and genomic DNA isolation and qPCR

Cultured cells were washed with ice cold PBS and lysed on the place with Trizol^TM^. RNA was isolated using the DirectZol^TM^ RNA Microprep Kit (Zymo Research, R2062) with on-column DNase digestion followed by cDNA synthesis using the Transcriptor First Strand cDNA synthesis kit (Roche Molecular Diagnostics, 04379012001) using random primers. Genomic DNA for genotyping was isolated from mouse tail snips with lysis buffer containing Proteinase K (Sigma-Aldrich, P2308). qPCR was performed using GoTaq qPCR Master Mix (Promega, A6002). The input RNA amount was normalized across all samples and *18S rRNA* or *HtatsF1* was used for normalization of qPCR across samples. Primer sequences are listed in Table S3. All primers were synthesized as custom DNA oligos from Integrated DNA technologies (IDT).

### Single cell RNAsequencing and analysis

Cells for single cell RNAseq were obtained from hind limb muscles of 2-3 moth-old *Pax7^tdT^:Flk1^GFP^* mice following enzymatic digestion as previously described (Liu et al., 2015). Dead cells were excluded from the analysis using ZombieNIR (Biolegends, 423105). TdTomato(+) and GFP(+) cells were sorted individually and then 20% of GFP(+) cells were spiked into 80% tdTomato(+). We loaded ∼5,000 cells into 1 channel of the Chromium system for each of these samples and prepared libraries according to the manufacturer’s protocol using version 2.0 chemistry (10x Genomics). Following capture and lysis, we synthesized cDNA and amplified for 12 cycles as per the manufacturer’s protocol (10x Genomics). The amplified cDNA was used to construct Illumina sequencing libraries that were each sequenced with ∼300K read/cell on one lane of an Illumina HiSeq 2500 machine. We used Cell Ranger 3.1 (10X Genomics) to process raw sequencing data. For A custom genome was constructed to include *eGFP-SV40*, *tdTomato-WPRE-BGHPolyA* and *Pax7-IRES-CreERT2* transgenes. Detailed step-by-step instructions can be found at https://github.com/verma014/10XCustomRef. We carried out analyses of the filtered data using Seurat suite version 3.0 (Stuart et al., 2019) in R studio (RStudio Team, 2020). For cell imputation, we utilized ALRA through the Seurat wrapper with default settings(Linderman et al., 2018). Additional scRNAseq datasets were obtained from GEO and analyzed using the same method as listed above. A myogenic score was calculated based on the expression of *Myog*, *Pax7*, *Myod1* and *Myf5*. Step-by-step instructions for the analysis can be found on https://github.com/verma014/10XCustomRef.

### Background Subtraction

10x Genomics scRNAseq platform uses many more droplets than cells and so following a run, there are many droplets that do not have cells. These droplets still get sequenced with some of the RNA that is in the solution. This floating RNA can be used to estimate the “background” in each droplet. A better description of this can be found by the developers of ‘SoupX’ (Young and Behjati, 2020). Since *Cdh5* expression has previously been verified in MuSCs using RNAscope, we were able to use it as a positive control to remove the background or “soup” from our data. If *Cdh5* is absent from MuSC, we know that the background subtraction was too aggressive and that subtracting the Soup is not reliable in our case. In addition, we know certain genes that are considered to be specific for MuSCs, muscle ECs or muscle fibers based on the bulk RNAseq (Verma et al., 2018). The top 5 genes that are specific to these population (and also detected by 10x) were selected and used to show the background in our data set was 14.40% and 13.89% for the D0 and D3 dataset, respectively. The step-by-step instructions can be found on https://github.com/verma014/10XCustomRef.

### Bulk RNAseq and Microarray Analysis

FASTQ files were downloaded from SRA using SRA-toolkit. Sequences were trimmed using trimmomatic to remove adapter contamination and low-quality reads. Trimmed sequences were mapped to mouse mm10 using Hisat2 (Pertea et al., 2016). Transcript assembly was performed using StringTie (Pertea et al., 2016). Cell type specificity was determined as previously described (Verma et al., 2018). Microarray analysis was performed using the Affymetrix Transcriptome Analysis Console (TAC). Samples in each experiment were RNA normalized and the expression was acquired using the Affeymetrix Expression analysis console with gene level expression. Heatmaps were generated in Prism 9 (Graphpad, La Jolla, CA). The code for generating each graph is listed in the following table, along with the link to the data in tabular format.

### Quantification and Statistical Analysis

Statistical analysis was performed using Prism 9 (Graphpad, La Jolla, CA) or RStudio (RStudio Team, 2020). For comparison between two groups, an unpaired T-test was used. For comparison between multiple groups, a one-way ANOVA was used with multiple comparisons to the control. Distributions were compared using a chi-squared test. Graphing of the data was performed using Prism 9. Vector diagrams were modified using Graphic (Autodesk). All values are means ±SEM unless noted otherwise. * indicates p < 0.05, ** indicates p < 0.01, *** indicates p < 0.001.

## Data and Software Availability

All the data was obtained from NCBI GEO. Microarrays of mouse MuSCs were obtained from GSE3483 (Fukada et al., 2007). scRNAseq of MuSCs and muscle ECs was performed in this study (GSE129057). scRNAseq of whole muscle was obtained from GSE143437 (De Micheli et al., 2020). Bulk RNAseq of MuSCs, ECs and single muscle fibers was obtained from GSE108739 (Verma et al., 2018) and GSE64379 (Ryall et al., 2015). Bulk RNAseq of TU-tagged RNA of MuSCs was obtianed from GSE97399 (van Velthoven et al., 2017). Bulk RNAseq of fixed and unfixed MuSCs was obtained from GSE113631 (Yue et al., 2020). Exercise, ALS, DMD, BMD, FSHD GSE3307, Early DMD GSE465, *mdx* GSE466, GRMD GSE69040, Satellite cells GSE15155. All arrays were normalized to their respective controls. All arrays and RNAseq data are listed in Supplemental Table 1 (Table S1).

## Acknowledgements

We thank Minnesota Supercomputing Institute, University of Minnesota Imaging Center, University of Minnesota FACS Facility, and University of Minnesota Genomics Center for providing data for this paper. We also thank Jake Trask for critical reading of this paper. We thank Drs. Yosuke Mukouyama (National Institute of Health), Napoleone Ferrara (Genentech), Guo-Hua Fong (University of Connecticut) and Masatsugu Ema (Siga University of Medical Science) for providing *VEGFA^LoxP/LoxP^*, *Flt1^LoxP/LoxP^* and *Flk1-GFP* mice, respectively. This work was supported by NIHT32-GM008244 and NIHF30AR066454 to MV, NIAMS grant AR070231 to RSK, a fellowship of the Training Program in Stem Cell Research from the New York State Department of Health to A.P.K. (NYSTEM-C32561GG) and NIHR01AR062142, NIHR21AR070319, MDA Research Grant, and Regenerative Medicine Minnesota (RMM) Grant to AA.

