## Supplementary Data for "Endothelial cell signature in muscle stem cells validated by VEGFA-FLT1-AKT1 axis promoting survival of muscle stem cell"

**Supplementary Data (Verma, et al.)**

**Supplementary Figure Legends**

**Figure S1**

A), B), C) and D) MuSC and EC genes expressed in MuSC from various published repositories show a discrepancy in expression of EC related genes (GSE3483, Fukada et al, 2007; GSE64379, Ryall et al., 2015; GSE113631, Yue et al., 2020; GSE97399, van Velthoven et al., 2017)

E) Example of FACS isolation of $Flk1^{GFP}$(+) ECs and $Pax7^{tdT}$(+) MuSCs from $Pax7^{+/CreERT2}$:$R26R^{+/tdT}$:$Flk1^{+/GFP}$ mice.

F) Green (Green FL) and red (Red FL) fluorescence microscopic images of freshly isolated $Flk1^{GFP}$(+) ECs and $Pax7^{tdT}$(+) MuSCs from $Pax7^{+/CreERT2}$:$R26R^{+/tdT}$:$Flk1^{+/GFP}$ mice. Nuclei were counterstained with DAPI (blue). Scale bar indicates 50 µm.

G) EC fragment contamination with single muscle fiber (SMF) preparation. $Flk1^{GFP}$(+)ECs (arrow, green) from the $Pax7^{+/CreERT2}$:$R26R^{+/tdT}$:$Flk1^{+/GFP}$ mice can be found enwrapping the muscle fibers when single muscle fibers (SMFs) were isolated, and that may result in EC gene expression in muscle fiber preparation. Arrowheads indicate $Pax7^{tdT}$(+)satellite cells (red). Nuclei were counterstained with DAPI (blue). Scale bar indicates 10 µm.

H) FACS shows clear delineation between the SSC and FSC in quiescent vs. activated MuSCs isolated from day 3-injured muscle of $Pax7^{+/CreERT2}$:$R26R^{+/tdT}$:$Flk1^{+/GFP}$ mouse.

I) Quantification of quiescent MuSCs (QSCs) vs. activated MuSCs (ASCs) based only on the FSC and SSC from basal and regenerating muscle from *Pax7*$^{+/CreERT2}$:*R26R*$^{+/tdT}$:*Flk1*$^{+/GFP}$ mice.

J) Genes used to define different cell identities in the scRNAseq. Of note, quiescent MuSCs were supported by high expression of *Hes1* in myogenic cells, activated MuSCs were defined as myogenic cells with high expression of *Ccnd1* and myoblasts were defined as myogenic cells expressing *Myogenin* (*Myog*). Activated ECs were defined as the population having higher *Ccnd1* expression.

K) Background subtraction using SoupX maintains EC identity in the *tdTomato*(+) and *CreERT2*(+) myogenic cell clusters as evident by preserved expression of EC genes (*eGFP* and *Cdh5*).

L) *Calcr*(+), *Myog*(+) and *Cdk1*(+) myogenic cell cluster from aggregated scRNAseq collection (GSE143437, De Michell et al., 2020) shows *Flt1* is expressed diffusely throughout the whole myogenic population.

**Figure S2**

A) MuSC-derived myoblast proliferation as assessed by % EdU following 6 hours of induction with exogenous VEGFA is increased at 20 ng/ml but decreased at 100 ng/ml.

B) Representative images of myoblast differentiation for 1 day stained with MyHC following bFGF or VEGFA treatment. Nuclei were counterstained with DAPI (blue). Scale bar indicates 50 μm.

C) Quantification of experiment shown in panel A showing that fusion index (equal or greater than 2 nuclei per cell) following 3 days in the differentiation medium is reduced by exogenous 20 ng/ml bFGF but unchanged with 20 ng/ml VEGFA.

D) *Myh3* expression normalized by *Htatsf1* in myoblasts was unchanged with exogenous VEGFA following 3 days in the differentiation medium.

E) Representative image of myoblast stained with crystal violet treated with VEGFA following induction of apoptosis via UV light. Scale bar indicates 200 µm.

F) Quantification of experiment shown in panel E shows improved cell survival of myoblasts with 20 ng/ml of VEGFA following induction of apoptosis via UV light.

G) Example of Annexin V/PI staining used for quantification of apoptosis by FACS shows the transition from live cells to apoptotic cells and necrotic cells following thapsigargin induction.

H) Time course of thapsigargin-induced cell apoptosis shows that cell survival is reduced to ~50% at 24 hours following thapsigargin induction.

I) RT-qPCR shows deletion of the *Flt1* exon 3 in *MuSC-Flt1*$^{\Delta/\Delta}$ myoblasts while exons 1 and 2 are retained in *MuSC-Flt1*$^{+/+}$ myoblasts.

J) Anti-FLT1 antibody staining shows downregulation of FLT1 in *MuSC-Flt1*$^{\Delta/\Delta}$ myoblasts while FLT1 is detected in *MuSC-Flt1*$^{+/+}$ myoblasts. Nuclei were counterstained with DAPI (blue). Scale bar indicates 50 µm.

K) Myoblast proliferation as assessed by % EdU in *MuSC-Flt1*$^{\Delta/\Delta}$ cells was not significantly altered compared with *MuSC-Flt1*$^{+/+}$ cells.

L) Myogenic differentiation as assessed by MyHC staining in *MuSC-Flt1*$^{\Delta/\Delta}$ myoblasts was not significantly altered compared with *MuSC-Flt1*$^{+/+}$ myoblasts following 3 days in the differentiation medium.

M) Representative images of myoblast cultures following EdU exposure in growth medium stained for EdU (green) and MyHC (red). Nuclei were counterstained with DAPI (blue). Scale bar indicates 50 µm.

**Figure S3**

A) *Vegfa* is increased in *VEGFA*$^{+/Hyper}$ mouse muscle but does not lead to an increase in *Flk1,* soluble *Flt1* (*sFlt1*) or membrane-bound *Flt1* (*mFlt1*).

B) Representative H&E images from *VEGFA*$^{+/+}$ and *VEGFA*$^{+/Hyper}$ mouse TA muscle in 28 days post-injury. Scale bar indicates 50 µm

C) Fiber size distribution and D) mean feret's diameter of uninjured muscle from *VEGFA*$^{+/Hyper}$ mice show an increase in fiber diameter compared with the control following injury.

E) Anti-VEGFA antibody staining shows downregulation of VEGFA protein in *MuSC-VEGFA*$^{\Delta/\Delta}$ myoblasts while VEGFA is detected in *MuSC-VEGFA*$^{+/+}$ myoblasts. Pax7$^{tdT}$(+) cells was clealy detected only in *MuSC-VEGFA*$^{\Delta/\Delta}$ myoblasts but not in *MuSC-VEGFA*$^{+/+}$ myoblasts. Nuclei were counterstained with DAPI (blue). Scale bar indicates 50 µm.

F) Representative H&E and Oil Red O (showing fat infiltration as red color) images from regenerating TA muscle of *MuSC-VEGFA*$^{+/+}$ and *MuSC-VEGFA*$^{\Delta/\Delta}$ mice in 28 days post-injury. Scale bar indicates 50 µm

G) Fiber size distribution and H) average fiber diameters are decreased in *MuSC-VEGFA*$^{Δ/Δ}$ mouse TA muscle compared with the control from *VEGFA*$^{+/+}$ mouse TA muscle

I) Average Oil Red O(+) fat infiltration from regenerating TA muscle of *MuSC-VEGFA*$^{+/+}$ and *MuSC-VEGFA*$^{Δ/Δ}$ mice in 28 days post-injury.

J) Representative H&E images from *MuSC-Flt1*$^{+/+}$ and *MuSC-Flt1*$^{Δ/Δ}$ mouse TA in 7 and 14-days post-injury. Scale bar indicates 50 μm

K) Fiber size distribution and L) mean feret's diameter are decreased 7 days post-injury in *MuSC-Flt1*$^{Δ/Δ}$ mouse TA muscle compared with the control from *MuSC-Flt1*$^{+/+}$ mouse TA muscle.

M) Fiber size distribution and N) average fiber diameter are decreased 14 days post-injury in *MuSC-Flt1*$^{Δ/Δ}$ mouse TA muscle compared with the control from *MuSC-Flt1*$^{+/+}$ mouse TA muscle

**Figure S4**

A) Heatmap of microarray data profiling *Vegfa* ligands and receptors in MuSC cultures (left), whole muscle in animal models (middle) and huma patient biopsy samples (right).

B) Body mass is preserved in *mdx:MuSC-Flt1*$^{Δ/Δ}$ compared with the control *mdx:MuSC-Flt1*$^{+/+}$ mice at both 3 and 12 months of age.

C) Fiber size distribution of *mdx:MuSC-Flt1*$^{Δ/Δ}$ compared with the control *mdx:MuSC-Flt1*$^{+/+}$ mouse TA muscle shows a small decrease in fiber size.

D) Body mass is preserved in *mdx*:*VEGFA*$^{+/Hyper}$ compared with the control *mdx*:*VEGFA*$^{+/+}$ mice at 12 months of age.

E) Fiber size distribution of *mdx*:*VEGFA*$^{+/Hyper}$ compared with the control *mdx*:*VEGFA*$^{+/+}$ mice TA and diaphragm (DM) muscle shows a small increase in fiber size.

### Supplemental Figure 1

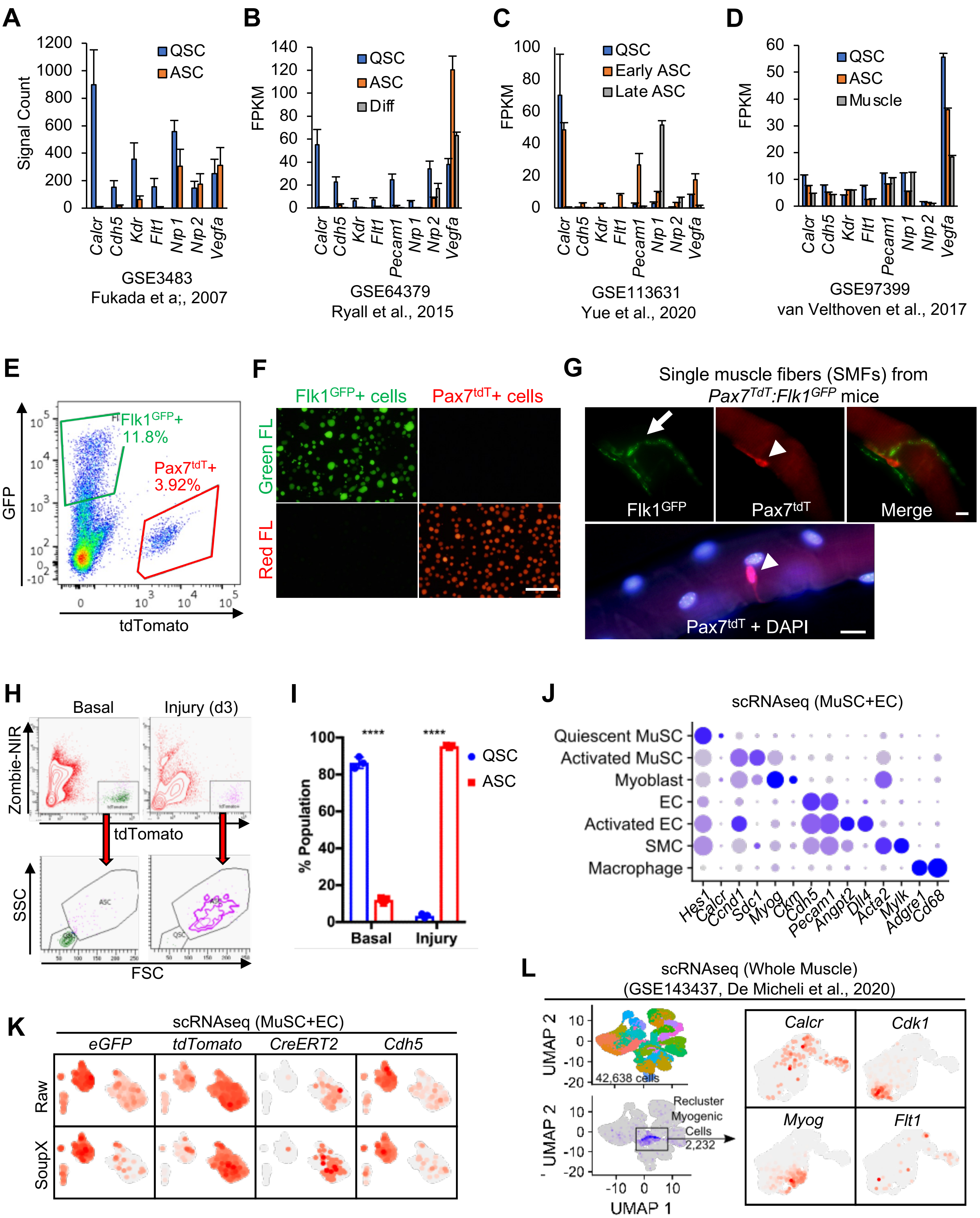

### Supplemental Figure 2

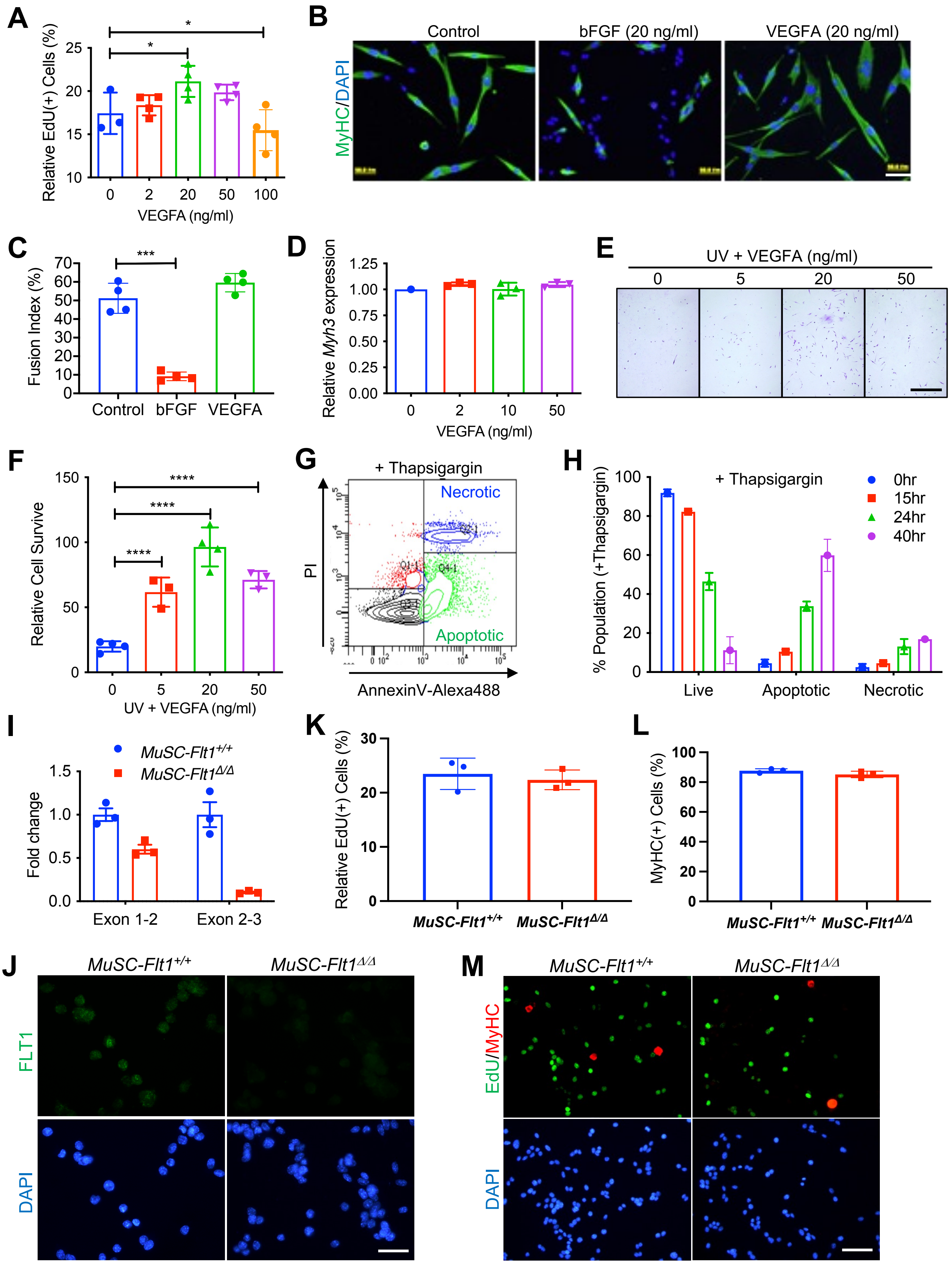

### Supplemental Figure 3

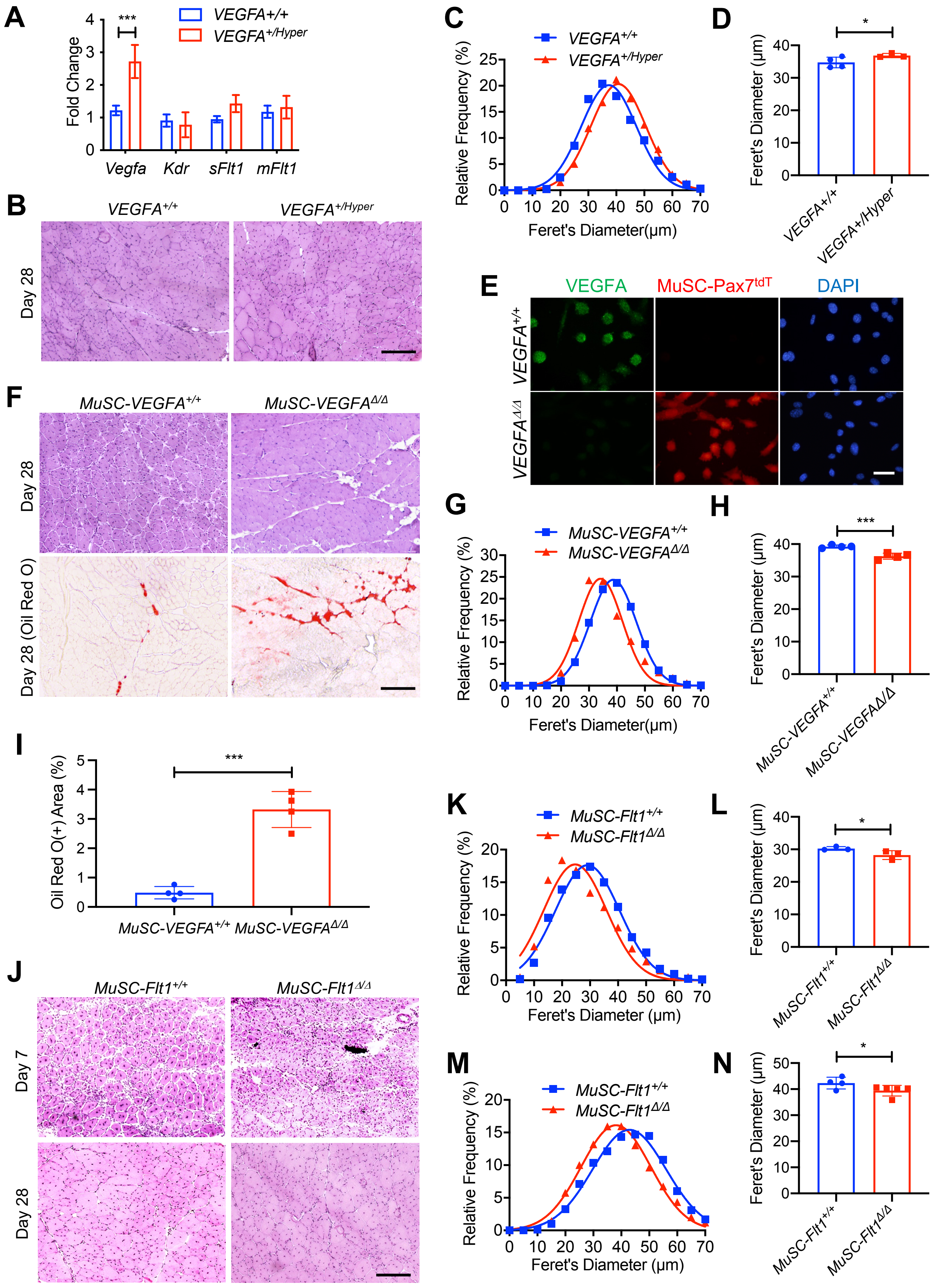

### Supplemental Figure 4

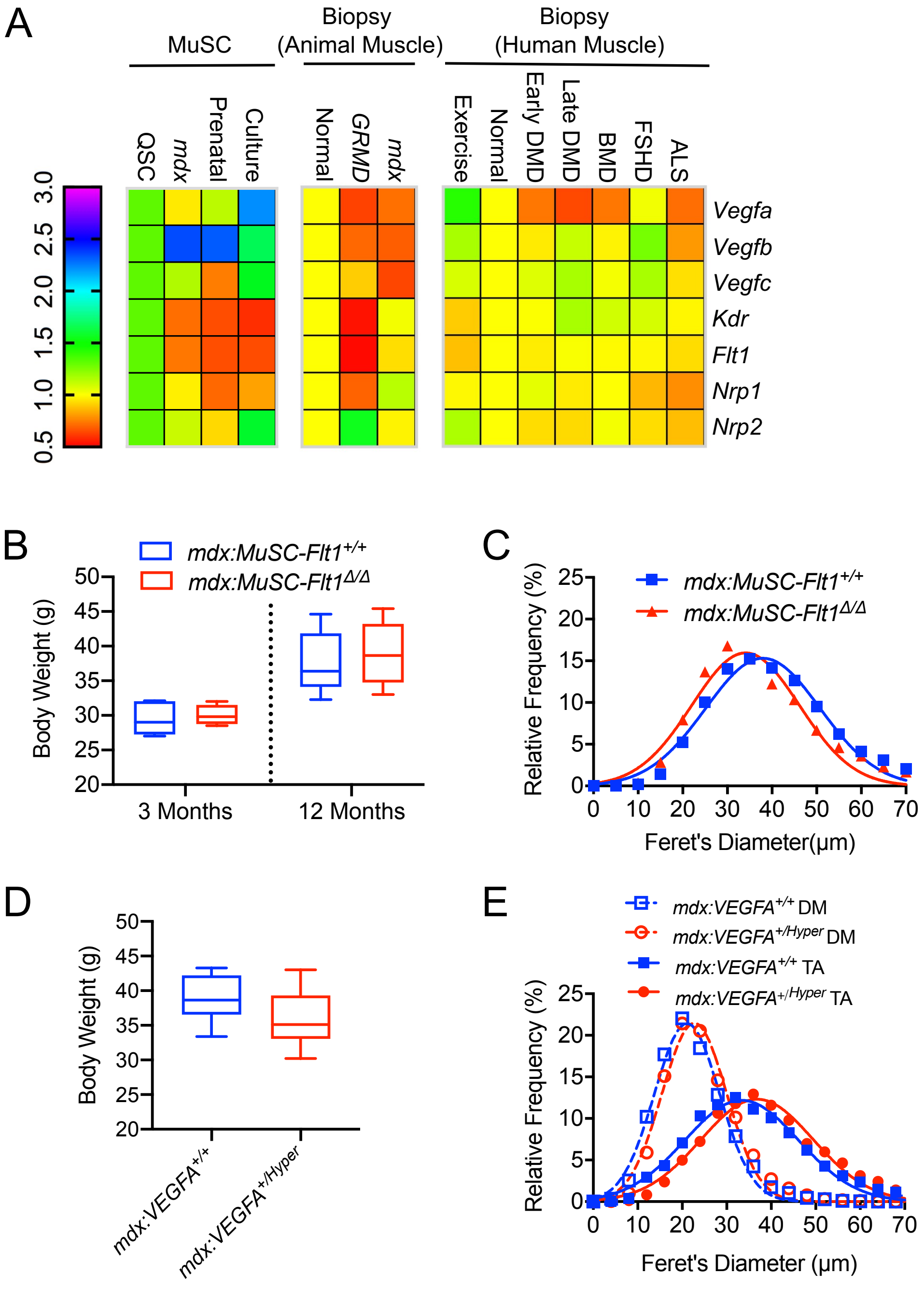

### Table S1

| Table S1. Gene Expression Omnibus (GEO) | | |
|---|---|---|
| **Bulk RNAseq, scRNAseq or Microarrays** | **Repository** | **Reference** |
| scRNAseq of MuSC and muscle ECs | GSE129057 | Current manuscript |
| scRNAseq of whole muscle | GSE143437 | De Micheli et al., 2020 |
| Bulk RNAseq of MuSCs, ECs and single muscle fibers | GSE108739 | Verma et al., 2018 |
| Bulk RNAseq of MuSCs | GSE64379 | Ryall et al., 2015 |
| Bulk RNAseq of TU-tagged RNA of MuSCs | GSE97399 | van Velthoven et al., 2017 |
| Bulk RNAseq of fixed and unfixed MuSCs | GSE113631 | Yue et al., 2020 |
| Microarrays of mouse MuSCs | GSE3483 | Fukada et al., 2007 |
| Microarrays of human muscle from exercise, ALS, DMD, BMD and FSHD | GSE3307 | Dadgar et al., 2014 |
| Microarrays of early human DMD muscle | GSE465 | Chen et al, 2000 |
| Microarrays of *mdx* mouse muscle | GSE466 | Tseng et al., 2002 |
| Microarrays of DMD dog muscle | GSE69040 | Vieira et al., 2017 |
| Microarrays of *mdx* mouse MuSCs | GSE15155 | Pallafacchina et al., 2010 |

### Table S2

| Mouse model | Treatment | Figures | Vascular change | Muscle pathology | Significant functional outcome |
|---|---|---|---|---|---|
| *VEGFA$^{+/Hyper}$* | N/A | 3, S3 | Increased capillary density | Improved muscle pathology | Not evaluated |
| *Pax7$^{CreERT2}$:VEGFA$^{LoxP/LoxP}$* (*MuSC-VEGFA$^{\Delta/\Delta}$*) | TMX | 3, S3 | Increased capillary density | Worse muscle pathology | Not evaluated |
| *Pax7$^{CreERT2}$:Flt1$^{LoxP/LoxP}$* (*MuSC-Flt1$^{\Delta/\Delta}$*) | TMX | 3, S3 | Not evaluated | Worse muscle pathology | Not evaluated |
| *mdx:Pax7$^{CreERT2}$:Flt1$^{LoxP/LoxP}$* (*mdx:MuSC-Flt1$^{\Delta/\Delta}$*) | TMX | 4, S4 | Increased capillary density | Worse muscle pathology | Reduced grip strength |
| *mdx:VEGFA$^{+/Hyper}$* | N/A | 4, S4 | Increased capillary density | Improved muscle pathology | Increased grip strength |

**Table S2. Summary of mice used for this paper**

### Table S3

| Table S3 DNA primer sequences | | | | | |
|---|---|---|---|---|---|
| **Gene** | **Forward** | **Sequence** | **Reverse** | **Sequence** | **Product (bp)** |
| **Genotyping** | | | | | |
| *Pax7$^{CreERT2}$* | Common | GCTGCTGTTGATTACCTGGC | WT reverse | CTGCACTGAGACAGGACCG | 419 (WT) |
| *Pax7$^{CreERT2}$* | Common | GCTGCTGTTGATTACCTGGC | MUT reverse | CAAAAGACGGCAATATGGTG | 235 (MUT) |
| *R26$^{tdT}$* | 9020 (WT) | AAGGGAGCTGCAGTGGAGTA | 9021 (WT) | CCGAAAATCTGTGGGAAGTC | 297 (WT) |
| *R26$^{tdT}$* | 9103 (MUT) | GGCATTAAAGCAGCGTATCC | 9105 (MUT) | CTGTTCCTGTACGGCATGG | 196 (MUT) |
| *Flk1$^{GFP}$* | GFP-F1 | AGCAGCACGACTTCTTCAAGTCCG | GFP-R1 | CTCCTTGAAGTCGATGCCCTTCAG | 161 |
| *VEGFA$^{Hyper}$* | 24286 (WT) | CCA TAG ATG TGA CAA GCC AAG | 25308 (Common) | GAC CGT GCT TGG TCA CCT | 254 (WT) |
| *VEGFA$^{Hyper}$* | 25307 (MUT) | ACC CGG GGA TCC TCT AGA AC | 25308 (Common) | GAC CGT GCT TGG TCA CCT | 199 (MUT) |
| *VEGFA$^{LoxP/LoxP}$* | muVEGF 419.F | CCTGGCCCTCAAGTACACCTT | muVEGF 567.R | TCCGTACGACGCATTTCTAG | 106 (WT) 148 (MUT) |
| *Flt1$^{LoxP/LoxP}$* | FlpeEX3F | CGCTTTTTGTCAGTCATCTTCA | FlpeInt3R | GAGAATGCACTGTGCTGAAGGA | 223 (WT) 641 (MUT) |
| *mdx$^{5cv}$* | 0981 | GAAGCTCCCAGAGACAAGTC | 0982 | TCATGAGCATGAAACTGTTCTT | 180 (WT, MUT) |
| **RT-qPCR** | | | | | |
| *Flt1* | Exon 1 | CTTGCTCACCATGGTCAGCTGCTG | Exon 2 | CACTTTTAACTTCGACCCTGAGCC | 103 (WT/KO) |
| *Flt1* | Exon 2 | GGCCAGACTCTCTTTCTCAAGTGC | Exon 3 | GCAGAATTGCCTGTTATCCCTCCC | 135 (WT) |
| *Flt1* | Flt1-F1 | GCAGAGCCAGGAACATATACACA | Flt1-R1 | GAGATCCGAGAGAAAATGGCCTTT | 103 |
| *Kdr (Flk1)* | Flk1-F1 | CAGTGGTACTGGCAGCTAGAAG | Flk1-R1 | ACAAGCATACGGGCTTGTTT | 66 |
| *Nrp1* | Nrp1-F1 | TCCTGGGAAACTGGTATATCTATGA | Nrp1-R1 | CATTCCAGAGCAAGGATAATCTG | 75 |
| *Nrp2* | Nrp2-F1 | ATGGCTGGACACCCAATTT | Nrp2-R1 | ATGGTTAGGAAGCGCAGGT | 67 |
| *Myh3* | Myh-F1 | CACCTGGAGAGGATGAAGAAGAA | Myh-R1 | AAGACTTGACTTTCACTTGGAGTTTATC | 298 |
| *Htatsf1* | Htatsf1-F1 | TTCGGAAGCTCCTTCTGTTT | Htatsf1-R1 | CCAGAGTCTGAATACAATGGTCA | 79 |
| *18S rRNA* | 18S-F1 | CGCACGGCCGGTACAGTGAAACTG | 18S-R1 | CACCCGTGGTCACCATGGTAGGCA | 343 |

### Table S4

| **Table S4 Antibodies** | | | | |
|---|---|---|---|---|
| **Antibodies** | | | | |
| **Name** | **Company** | **Catalogue #** | **Use case** | **Dilution** |
| FLK1-APC | eBioscience | 17-4321-81 | FACS | 1:100 |
| FLT1-APC | R&D systems | FAB4711A | FACS | 1:100 |
| NRP1-APC | R&D systems | FAB4711A | FACS | 1:100 |
| NRP2-APC | R&D systems | FAB22151A | FACS | 1:100 |
| IgG-APC | eBioscience | 17-4321-81 | FACS | 1:100 |
| PECAM1 | eBioscience | 14-0311-85 | IF | 1:200 |
| VE-Cadherin | eBioscience | 14-1441-82 | IF | 1:200 |
| VEGFA | Abcam | AB52917 | IF | 1:200 |
| FLK1 | eBioscience | 14-5821-82 | IF | 1:200 |
| FLT1 | R&D systems | MAB4711 | IF | 1:200 |
| NRP1 | Cell Signaling Technology | D62C6 | IF | 1:200 |
| NRP2 | Cell Signaling Technology | D39A5 | IF | 1:200 |
| MyHC | Developmental Study Hybridoma Bank (DSHB) | MF-20 | IF | 1:50 |
| MyoD | Santa Cruz Biotechnology | sc-304 | IF | 1:500 |
| Pax7 | Developmental Study Hybridoma Bank (DSHB) | Pax7 | IF | 1:20 |
| pAKT | Cell Signaling | 4060 | IF | 1:500 |
| Anti-Mouse Alexa Fluro 488 | ThermoFisher Scientific | A-21202 | IF | 1:1000 |
| Anti-Rabbit Alexa Fluro 488 | ThermoFisher Scientific | A-11008 | IF | 1:1000 |
| Anti-Rat Alexa Fluro 488 | ThermoFisher Scientific | A-21210 | IF | 1:1000 |
| Anti-Goat Alexa Fluro 488 | ThermoFisher Scientific | A-11078 | IF | 1:1000 |
